# A Single Laccase Acts as a Key Component of Environmental Sensing in a Broad Host Range Fungal Pathogen

**DOI:** 10.1101/2023.01.12.523834

**Authors:** Nathaniel M. Westrick, Eddie G. Dominguez, Christina M. Hull, Damon L. Smith, Mehdi Kabbage

**Affiliations:** Department of Plant Pathology, University of Wisconsin-Madison, Madison, WI, United States; United States Department of Agriculture – Agricultural Research Service, Madison, WI, United States; Department of Biomolecular Chemistry, University of Wisconsin-Madison, School of Medicine and Public Health, Madison, WI, United States; Department of Medical Microbiology and Immunology, University of Wisconsin-Madison, School of Medicine and Public Health, Madison, WI, United States

## Abstract

Secreted laccases are important enzymes on an ecological scale for their role in mediating plant-fungal interactions, but their function in fungal pathogenesis has yet to be elucidated. Ascomycete laccases have been primarily associated with cell wall melanin deposition, and laccase mutants in ascomycete species often demonstrate reduced pigmentation. In this study, a putatively secreted laccase, *Sslac2*, was characterized from the broad host-range plant pathogen *Sclerotinia sclerotiorum*, which is largely unpigmented and is not dependent on melanogenesis for plant infection. Of the seven putative laccases in the *S. sclerotiorum* genome, *Sslac2* was the only one found to be highly upregulated during pathogenesis of soybeans and was additionally found to be induced during growth on solid surfaces. Gene knockouts of *Sslac2* demonstrate wide ranging developmental defects, including abolished sclerotial formation, and are functionally non-pathogenic on unwounded tissue. While these mutants demonstrated a clear radial growth defect, enhanced growth was observed in liquid culture, likely due to altered hydrophobicity and thigmotropic responsiveness. *Sslac2* mutants were also unable to respond to a host environment, and accordingly unable to differentiate penetration structures, respond appropriately to chemical stress, or produce the key virulence factor oxalic acid. Transmission and scanning electron microscopy of WT and mutant strains show apparent differences in extracellular matrix structure that may explain the inability of the mutant to respond to environmental cues. Targeting *Sslac2* using host-induced gene silencing significantly improved resistance to *S. sclerotiorum*, suggesting that fungal laccases could be a valuable target of disease control. Collectively, we identified a laccase critical to the development and virulence of the broad host-range pathogen *S. sclerotiorum* and propose a potentially novel role for fungal laccases in modulating environmental sensing.

## Introduction

Laccases are a broadly conserved class of multicopper oxidase (MCO), which are known for their capacity to oxidize otherwise recalcitrant phenolic compounds, both directly and indirectly through the activity of mediators (Morozova et al. 2014). While these enzymes have garnered industrial interest in recent years as drivers of xenobiotic bioremediation, they play a pivotal role in development across a range of species and can be found in the genomes of plants, animals, fungi, oomycetes, and bacteria (Arregui et al. 2019; Feng et al. 2015; Morozova et al. 2014). One of their most important roles in natural ecosystems is to mediate the interplay between lignin production and degradation in interactions between plants and fungi. Higher plants utilize laccases in the biosynthesis and polymerization of lignin, which acts as a structural compound facilitating plant development and helps to defend plant tissue in response to pathogenesis (Arregui et al. 2019). Intriguingly, many wood rot fungi similarly use secreted laccases in the degradation of lignin, in which terminal phenolic lignin is directly oxidized by laccases and non-phenolic lignin is degraded through the activity of mediators, which are oxidized by the enzyme (Ten Have and Teunissen 2001). This interplay between production and degradation is critical to carbon cycling in the environment and the maintenance of healthy soils.

Unlike wood rot fungi, which are almost exclusively basidiomycetes, laccase activity within ascomycete fungi is typically associated with melanin/pigment deposition on fungal tissue, with most laccase knockout mutants within the phylum demonstrating reduced pigmentation (Fang et al. 2010; Lin et al. 2012; Lu 2017; Ma et al. 2017; Saitoh, Izumitsu, Morita, Shimizu, et al. 2010; Upadhyay et al. 2016; Wei et al. 2017). The best studied examples of such laccases are *Abr2* and *ya*, from *Aspergillus fumigatus* and *A. nidulans*, respectively, which function in melanization through the polymerization of phenolic monomers into mature melanin in fungal cell walls (Upadhyay et al. 2016). Intriguingly, the effect on fungal development and growth within these various knockouts is highly variable, likely due to the expanded repertoire of laccases and possible functional redundancy (Feng et al. 2015). This redundancy has been suggested by the broad substrate overlap observed within individual species’ laccase repertoires and demonstrated in mutants of the *Cochliobolus heterostrophus* laccase *ChMCO1*, in which mutant pigmentation defects could be complemented through the chemical induction of other laccases in *in-vitro* (Baldrian 2006; Saitoh, Izumitsu, Morita, Shimizu, et al. 2010). In the plant pathogen *Fusarium oxysporum* f. sp. *lycopersici*, multiple distinct laccases were knocked out, and while some differences in sensitivity to oxidative/chemical stress could be measured, no clear changes to development or virulence were observed (Cordoba Cañero and Roncero 2008).

In some cases, however, roles for specific laccase genes have been noted, such as in *Colletotrichum gloeosporioides, Colletotrichum orbiculare*, and *Setosphaeria turcica* where individual laccase gene knockouts result in reduced virulence on host plants (Lin et al. 2012; Ma et al. 2017; Wei et al. 2017). These laccases and the laccase *Mlac1* from the entomopathogenic fungus *Metarhizium anisopliae* are also deficient in appressorium formation (Fang et al. 2010; Lin et al. 2012; Ma et al. 2017; Wei et al. 2017). Given that these laccases are almost always associated with pigmentation in ascomycetes, it has broadly been assumed that a lack of melanin/pigment deposition is driving the wide variety of phenotypes observed in knockouts, although a causal relationship has never been confirmed (Lin et al. 2012; Lu 2017; Wei et al. 2017).

*Sclerotinia sclerotiorum* is a broad-host-range pathogen of dicotyledonous plants and is distinct from many of the previously mentioned fungi in that its melanin biosynthesis pathway has been relatively well studied. Three genes, encoding a scytalone dehydratase (*SCD1*), a trihydroxynaphthalene reductase (*THR1*), and a polyketide synthase (*PKS13*), with putative upstream roles in melanin biosynthesis have been knocked out and characterized (Li et al. 2018; Y. Liang et al. 2018). Although mutants were deficient in melanization of either sclerotia (*SCD1* and *THR1*) or compound appressoria (*PKS13*), no changes in virulence were observed, suggesting that melanin deposition does not significantly alter *S. sclerotiorum* pathogenicity (Li et al. 2018; Y. Liang et al. 2018). This is mirrored in the closely related species *Botrytis cinerea* in which melanogenic genes are dispensable for proper growth and virulence (Schumacher 2016). Of note, the *B. cinerea* laccase *Bclcc2* is broadly believed to be involved in the oxidation of environmental antimicrobials (Schouten et al. 2002, 2008). In this study we identify and characterize a *S. sclerotiorum* laccase gene, *Sslac2*, which is highly upregulated during infection of soybean and is the putative ortholog of the *B. cinerea* laccase *Bclcc2*. Contrary to *Bclcc2*, which has no clear role in development or virulence, *Sslac2* operates as a global regulator of environmental sensing, with *ΔSslac2* strains demonstrating clear alterations to every stage of development, infection, and response to environmental cues. This is unlike other fungal systems, such as *C. orbiculare*, where orthologous laccases between related species have been shown to be functionally interchangeable (Lin et al. 2012). In this study we also discuss the developmental and virulence phenotypes seen in *ΔSslac2* mutants, demonstrate the potential for targeting fungal laccases to achieve increased resistance in host plants, and consider the broader context of laccases within fungal and pathogen biology.

## Results

### Sslac2 is the primary laccase expressed during pathogenesis

Some fungal laccases play a role in plant pathogenicity, but fungi typically maintain multiple laccases within their genomes, so an evaluation of the *S. sclerotiorum* laccase repertoire was performed (Feng et al. 2015). Genomic analysis identified 7 putative laccases within the *S. sclerotiorum* genome, all of which contained predicted secretion signal domains, but which varied in length and cupredoxin domain architecture (Fig. 1A). While enzyme secretion is typically mediated by the presence of a signal peptide on the N-terminus of a protein, the C-terminus of fungal laccases is additionally processed during secretion and is considered critical to laccase activity (Andberg et al. 2009). The canonical motif mediating this activity is a C-terminal DSGL, and *Sslac2-6* contain a conserved DSGx motif in their C-terminus, although this is apparently missing in *Sslac1* and *Sslac7* (Fig. S1)(Andberg et al. 2009). To determine the laccases most important during *S. sclerotiorum* pathogenesis, a transcriptomic dataset generated from *S. sclerotiorum* infection of soybean at 24, 48, and 96 hours post inoculation (HPI) was analyzed to identify which homologues were upregulated *in-planta* compared to an *in-vitro* culture control (Westrick et al. 2019). Surprisingly, only *Sslac2* appeared to be highly upregulated during infection, particularly in the early stages of disease development (Fig. 1B).

**Figure 1.**
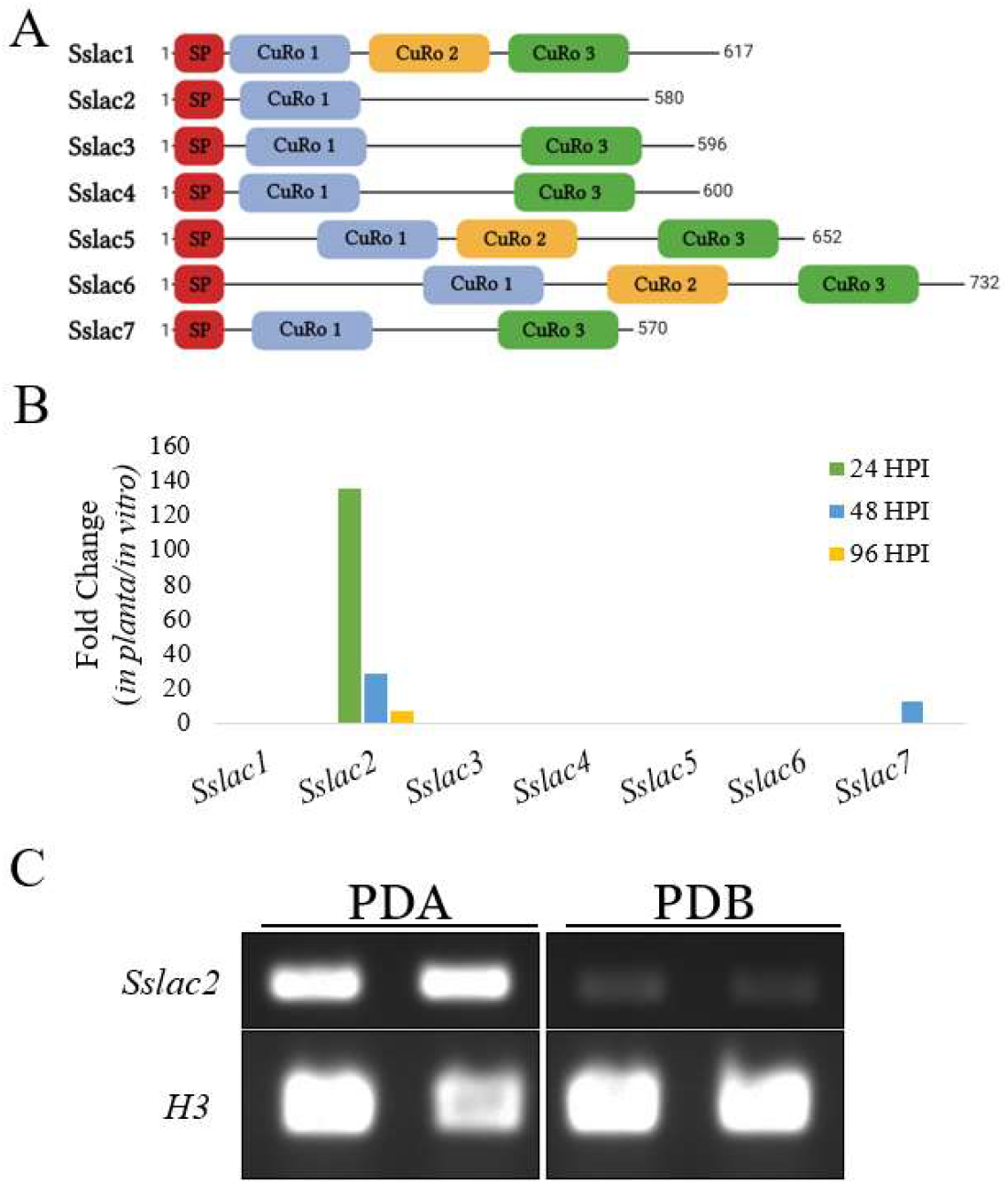
Schematic organization and expression of laccase genes in the genome of *S. sclerotiorum*. A) Length and domain architecture of the seven putative laccases in the *S. sclerotiorum* genome. SP refers to a predicted secretion signal peptide. CuRo 1, 2, and 3 correspond to the three cupredoxin domains characterized in *Melanocarpus albomyces* (cd13854, cd13901, and cd13880, respectively). B) Relative transcript fold change of the seven laccases identified in the *S. sclerotiorum* genome during infection of soybean when compared to *in vitro* culture growth. Fold change values are from the transcriptomic analysis performed in Westrick et al., 2019. C) RT-PCR of *Sslac2* and Histone 3 (*H3*) expression when grown on Potato Dextrose Agar (PDA) or in Potato Dextrose Broth (PDB).

Because the onset of infection requires the pathogen to interact with plant surfaces, it was considered whether any *S. sclerotiorum* laccases were distinctly upregulated on solid surfaces. To examine this, a separate transcriptomic dataset was analyzed from *S. sclerotiorum* grown on potato dextrose agar (PDA) or in potato dextrose broth (PDB), as these two mediums are nearly chemically identical outside of surface rigidity produced by the agar (Peyraud et al. 2019). Once again, *Sslac2* appeared to be the primary laccase upregulated (Fig. S2). This upregulation was additionally validated through RT-PCR (Fig. 1C). In toto, these data indicate that *Sslac2* is likely upregulated upon deposition of the pathogen on plant material.

### Laccase activity and sclerotial production are abolished in *ΔSslac2* knockout mutants

To evaluate the role of *Sslac2* in disease and development, a CRISPR-Cas9 assisted method was used to generate two independent *Sslac2* gene knockouts. Surprisingly, *ΔSslac2* strains completely lost the ability to produce sclerotia, and this was accompanied by an increase in the formation of arial hyphae (Fig. 2A). Laccase activity of these mutants was also evaluated by growing WT (1980) and mutant strains on media containing 0.2 mM 2,2’-azino-bis(3-ethylbenzothiazoline-6-sulfonic acid (ABTS) or tannic acid and evaluating pigment production, as oxidation by laccases is known to generate a violet/blue pigment from ABTS and a brown pigment from tannic acid (Ma et al. 2017; Schouten et al. 2002). These pigments were clearly observed in plates growing the WT (1980) strain but were completely absent in the mutants (Fig. 2B). All strains were additionally grown on ABTS and copper sulfate (CuSO4) in an attempt to complement this laccase deficiency through the activation of other laccase homologs, as CuSO4 has been shown to induce laccase activity in related species (Buddhika, Savocchia, and Steel 2021; Saitoh, Izumitsu, Morita, Shimizu, et al. 2010). While a minor increase in ABTS oxidation can be seen in the WT (1980) plates, addition of CuSO4 did not appear to increase laccase activity in either of the knockout mutants, suggesting that *Sslac2* is the primary laccase utilized by *S. sclerotiorum* during growth and development on solid surfaces (Fig. 2B).

**Figure 2.**
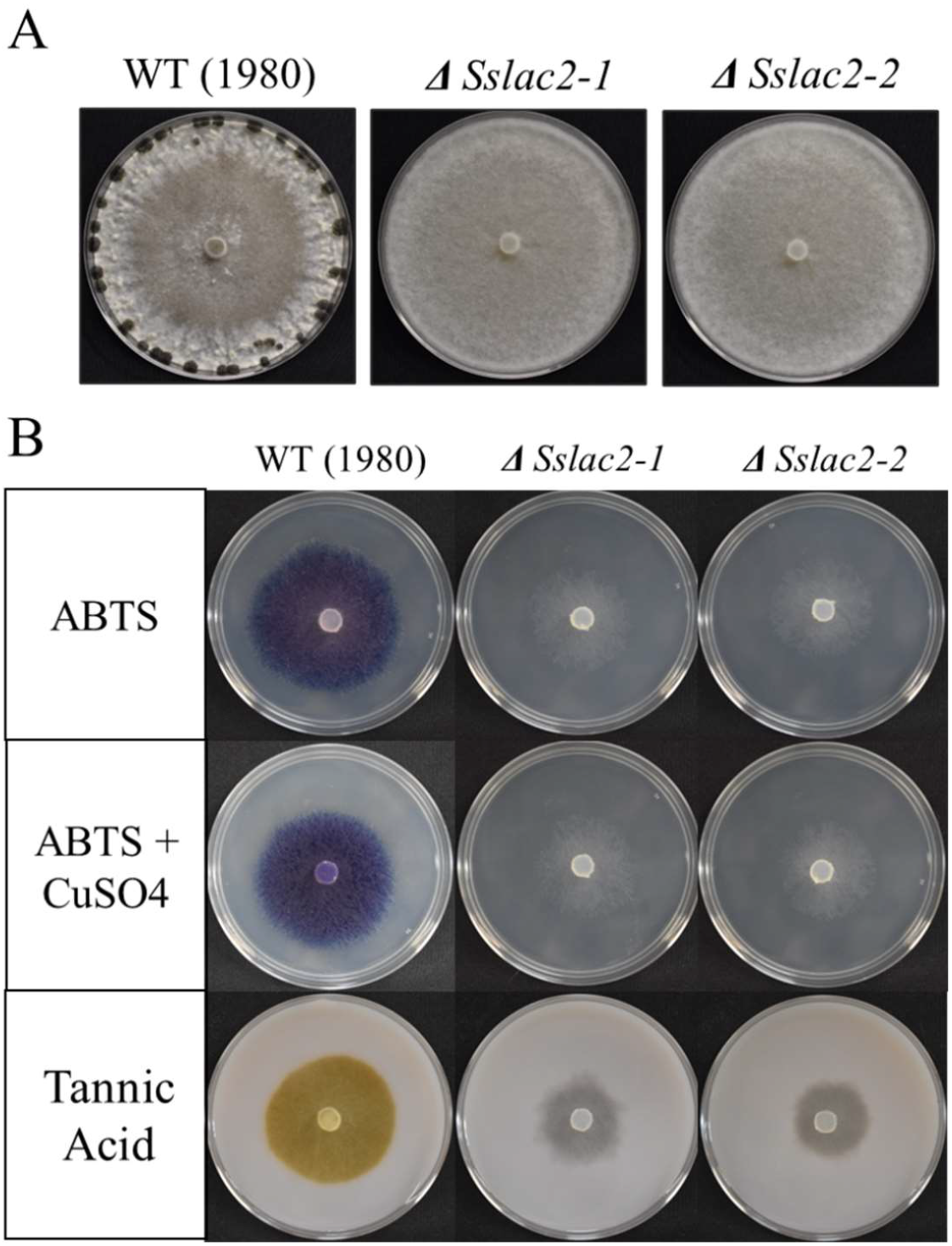
General phenotype and laccase activity of WT (1980) and laccase mutants. A) WT and mutant strains 2 weeks after inoculation of PDA plates. B) WT and mutant strains grown for 24 hours on PDA supplemented with 0.2 mM 2,2’-azino-bis(3-ethylbenzothiazoline-6-sulfonic acid (ABTS), 0.2 mM ABTS + 0.6 mM CuSO_4_, or 2.5 mg/ml tannic acid.

### *Sslac2* expression affects fungal tropism

Initial observations of *ΔSslac2* strains indicated a growth defect, as reduced radial growth could be observed on PDA (Fig. 3A). In contrast, when grown in liquid culture on PDB *ΔSslac2* mutants grew to a significantly higher mass than WT (1980), suggesting that the initial growth defect was specific to radial growth (Fig. 3B). A clear increase in arial hyphae was noted in older cultures of *ΔSslac2 (*Fig. 2A), so *ΔSslac2* and WT (1980) strains were grown for 2 weeks to assess whether the mutants were growing downward into the agar as well. At 2 weeks post inoculation (WPI), WT hyphae rested in a thin layer atop the agar, whereas *ΔSslac2* strains visibly penetrated the agar surface (Fig. 3C). Remarkably, when grown on split plates, *ΔSslac2* strains grew over the high barrier separating the plate sections and continued to grow into the neighboring chamber (Fig. S3). When confronted with such barriers, the WT strain typically enters dormancy and produces survival structures (Fig. S3). Thus, *ΔSslac2* may be unable to sense the environmental triggers leading to dormancy and the production of sclerotia.

**Figure 3.**
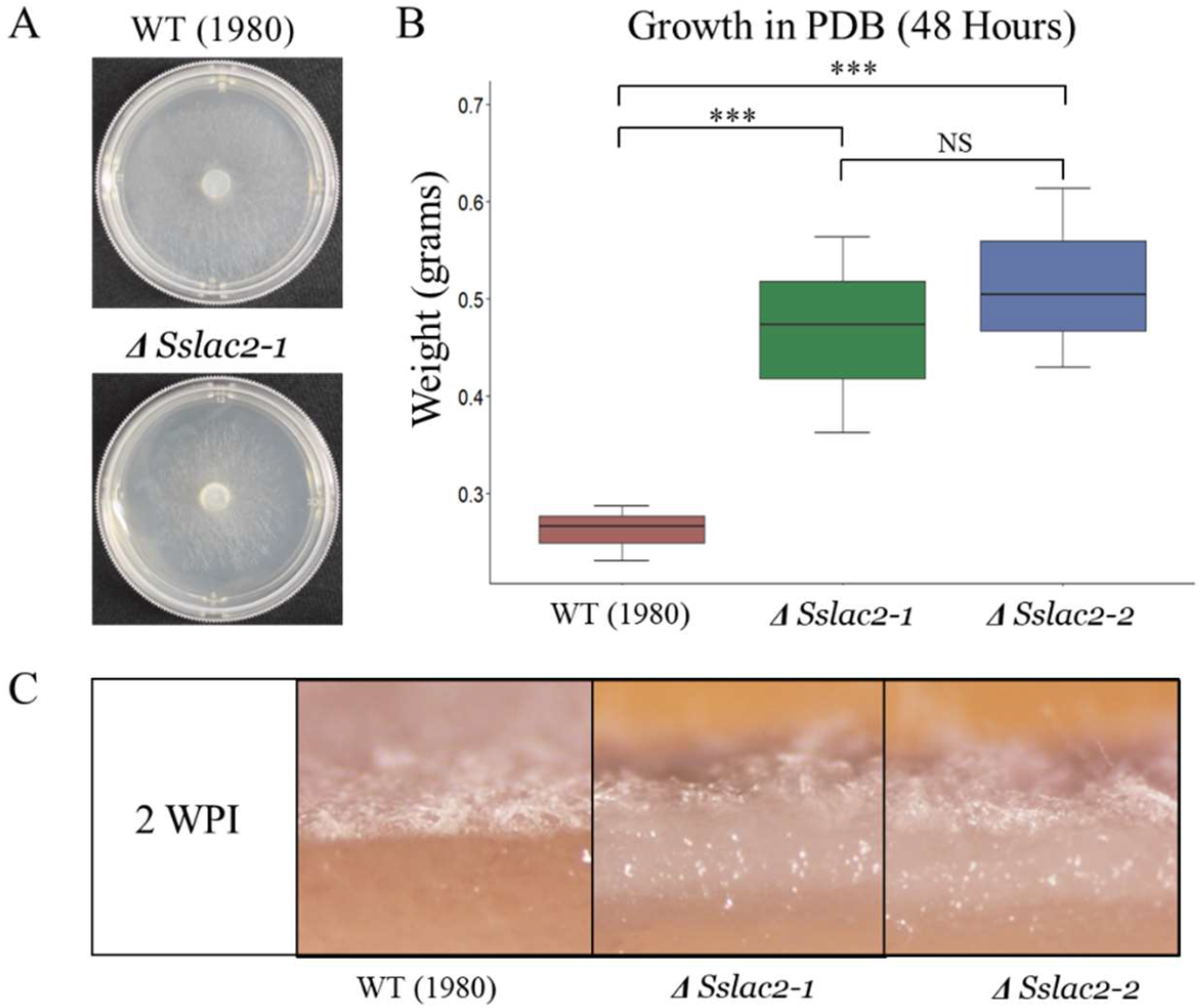
Phenotypes of WT and mutant strains grown on liquid and solid media. A) WT and *ΔSslac2-1* one day after inoculation on 60 mm PDA plates. B) Hyphal dry weight of WT and mutant strains grown in PDB for 48 hours. C) Cross section of PDA colonized with WT and mutant strains 2 weeks post inoculation (WPI). Statistical analysis utilized a Student’s t-test on three biological replicates of each strain (*<0.05, **<0.01, ***<0.001).

To assess if the agar penetration and slow radial growth phenotypes were caused by defects in directional growth or a failure to respond to dormancy triggers, all strains were grown on PDA with and without a top layer of sterile cellophane to halt agar penetration. While a significant defect in radial growth was measured on PDA, this defect disappeared when grown on cellophane (Fig 4). These data and the noted sclerotial phenotype in the mutants suggest that the mutant grows in a random, indiscriminate manner, whereas the WT prioritizes radial growth and is able to respond to environmental cues. This suggests that *Sslac2* plays a role in hyphal thigmotropism and the initiation of differentiation.

**Figure 4.**
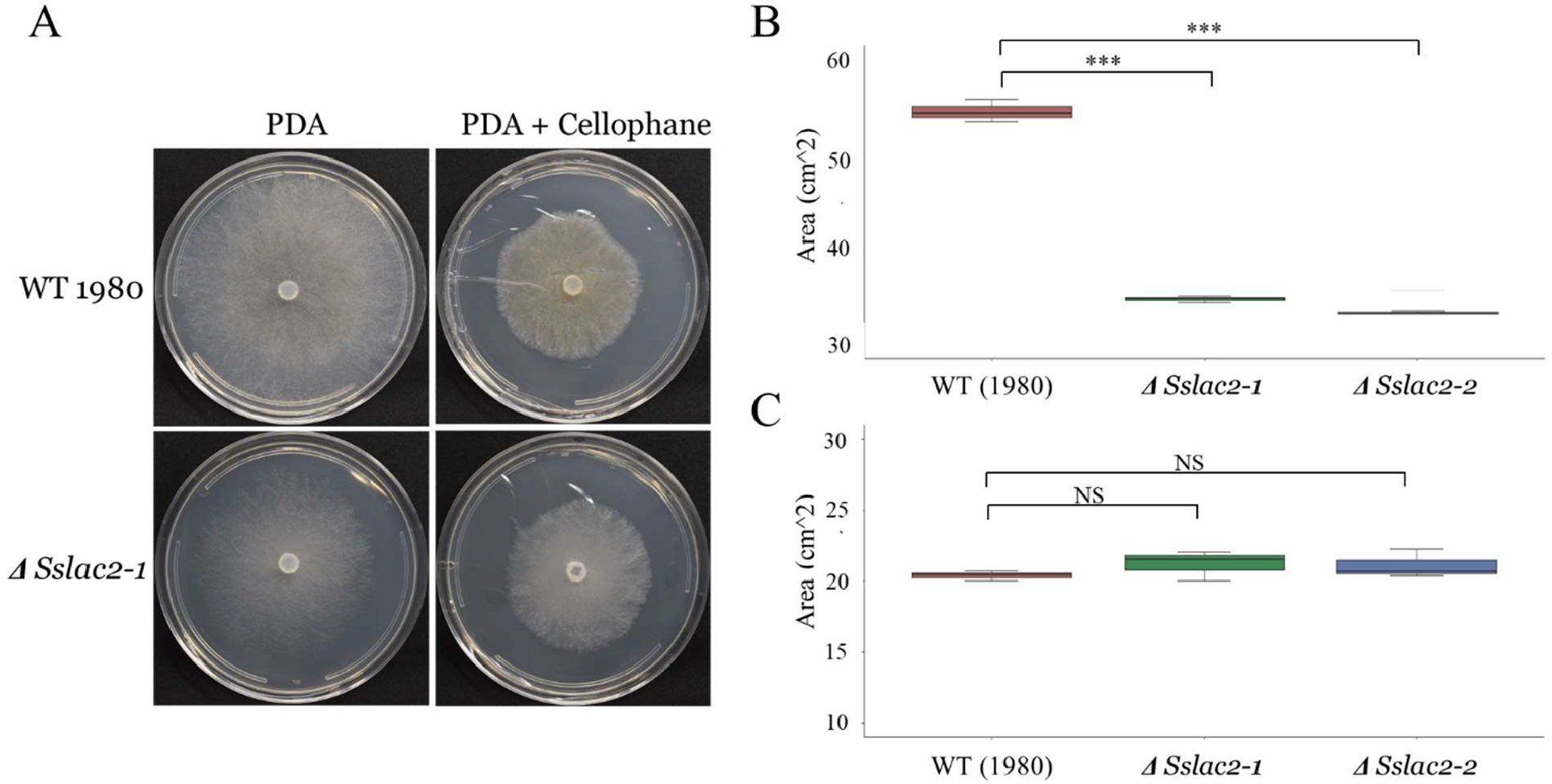
Growth of WT and mutants on PDA and cellophane covered PDA. A) WT and *ΔSslac2-1* strains 48 hours after inoculation. B) Quantification of colony area. Statistical analysis utilized a Student’s t-test on three biological replicates of each strain (*<0.05, **<0.01, ***<0.001).

### *Sslac2* is required for penetration structure formation and oxalic acid production

Because *Sslac2* is highly upregulated during the early stages of infection (Fig. 1B), we assessed the ability of *ΔSslac2* mutants to generate canonical compound appressoria (the penetration structures formed by *S. sclerotiorum* to infiltrate plant tissue). Strikingly, *ΔSslac2* mutants generated severely malformed compound appressoria in addition to generating far fewer when grown on glass slides, suggesting that *Sslac2* may be required for proper signaling and differentiation of penetration structures on host surfaces (Fig. 5A-B). Exogenously applied cAMP has been shown to rescue appressorium defects in G-protein-coupled receptor mutants; however, the addition of cAMP to the glass slides used for compound appressorium production did not rescue this defect for *ΔSslac2* mutants (data not shown)(Adachi and Hamer 1998).

**Figure 5.**
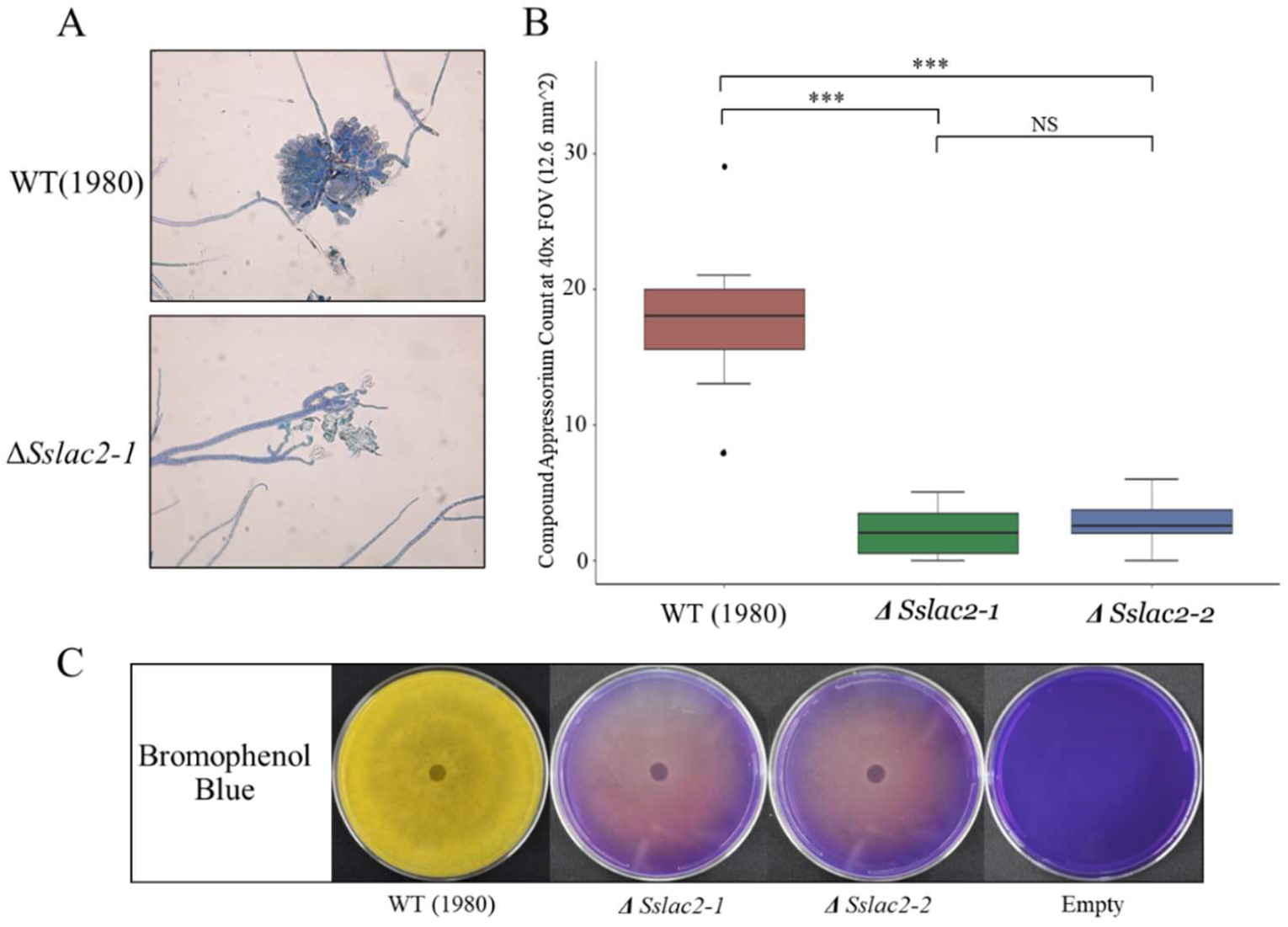
Compounds appressorium formation and oxalic acid secretion. A) Comparison of compound appressorium formation between WT (1980) and *ΔSslac2-1* strains. B) Quantification of canonical compound appressoria and attempted compound appressoria generated by the WT and mutant strains, respectively. C) Growth of the WT and mutant strains on bromophenol blue plates to qualitatively assess oxalic acid production. Statistical analysis utilized a Student’s t-test on three biological replicates of each strain (*<0.05, **<0.01, ***<0.001).

After penetrating into host tissue, *S. sclerotiorum* secretes copious amounts of oxalic acid, an organic acid that facilitates host tissue acidification as well as directly induces cell death and subverts defenses in the host (Mccaghey et al. 2018). Oxalic acid production was assessed using bromophenol blue plates, which shift from blue to yellow as the medium is acidified. While *ΔSslac2* mutants were capable of acidifying the media to a limited degree, they were clearly deficient relative to WT (1980) (Fig. 5C).

Given the importance of both penetration and oxalic acid production in *S. sclerotiorum* pathogenic development, it is unsurprising that these mutants were essentially non-pathogenic (Fig. 6A). This also highlights the importance of *Sslac2* as an environmental sensing component, leading to the production of both critical pathogenicity structures and virulence factors. To assess whether this defect in pathogenicity was due only to the loss of host penetration, WT and *ΔSslac2* strains were inoculated onto damaged soybean leaves, which provide direct access to the hosts tissues and have been shown in other systems to complement penetration structure defects (Fig. 6B). Additionally, *ΔSslac2* strains were compared with a *ΔSsOah1* strain deficient in oxalic acid production to assess the role that this organic acid may be playing in the *ΔSslac2* virulence defect (X. Liang et al. 2015). As expected, WT (1980) was capable of infecting both damaged and undamaged soybean leaves and *ΔSsOah1* colonization was limited to the area around the plant vasculature as previously reported (Xu et al. 2015). In contrast, even with damaged tissue, *ΔSslac2* strains were incapable of inducing lesions on soybean leaves, indicating that the previously observed loss of pathogenicity is not solely the result of defects in penetration and acidification (Fig. 6B). Because soybean is a moderately resistant host of *S. sclerotiorum*, we additionally tested *ΔSslac2* infection of the more susceptible host *Nicotiana benthamiana*. No lesions were observed on undamaged leaves (data not shown), but limited lesions could be seen on damaged leaves, albeit to a far lesser extent than either WT (1980) or *ΔSsOah1* (Fig. 6C).

**Figure 6.**
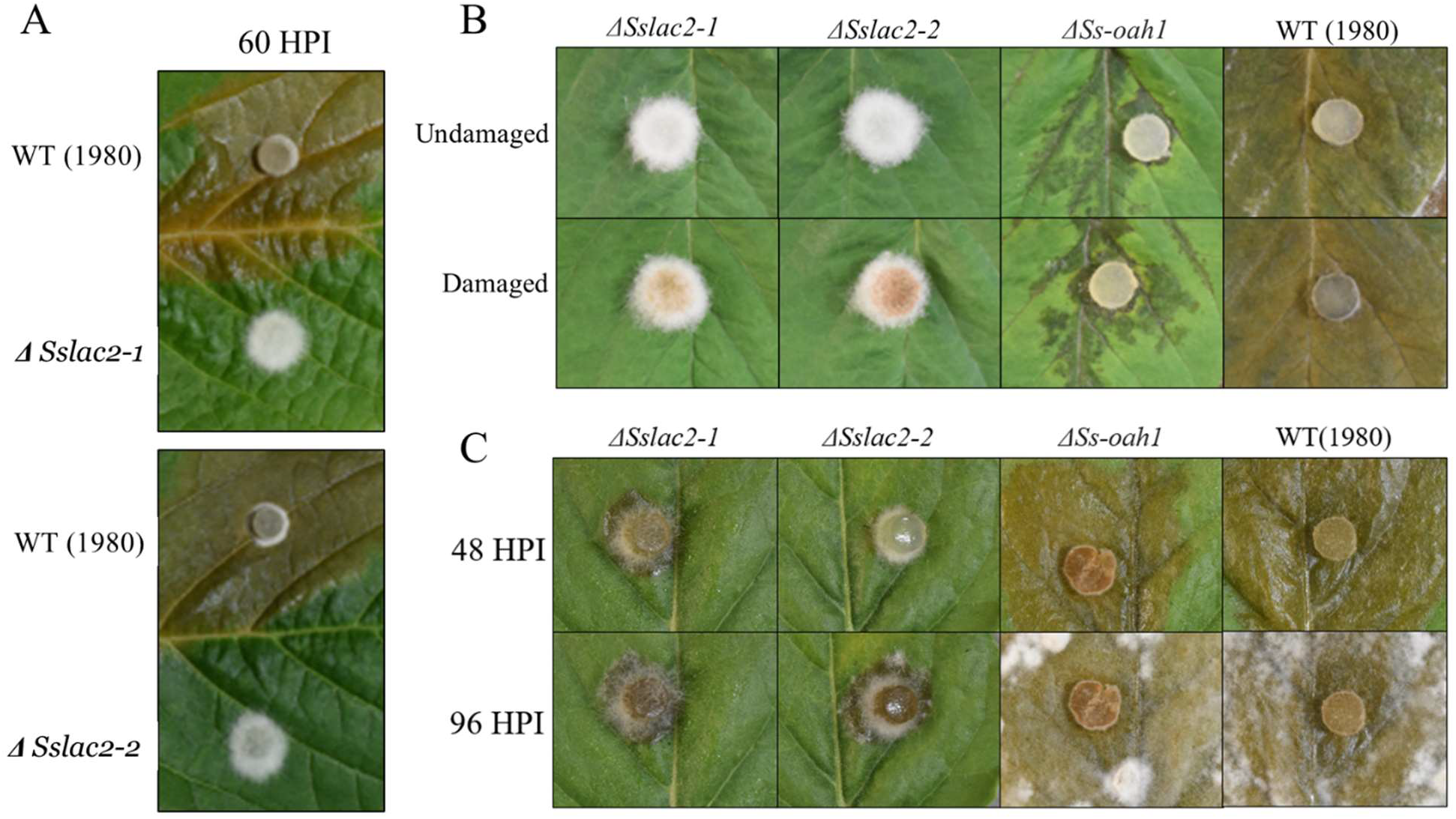
Virulence of WT and mutant strains. A) Comparison of WT (1980), *ΔSslac2-*1, and *ΔSslac2-2* during infection of soybean leaves. B) Infection of damaged and undamaged soybean tissue by WT (1980), *ΔSslac2-*1, *ΔSslac2-2*, and *ΔSs-oah1*. C) Infection of damaged *N. benthamiana* at 48 and 96 hours post infection (HPI) by WT (1980), *ΔSslac2-*1, *ΔSslac2-2*, and *ΔSs-oah1*.

While the *ΔSslac2* strain is able to marginally colonize wounded *N. benthamiana* leaves, overall, it is clear that this mutant’s virulence defect extends beyond penetration structures and oxalic acid production and likely involves the loss of other virulence factors or an inability to adequately respond to a hostile host environment.

### Transcriptional activity of *ΔSslac2* is less responsive to environmental factors

Many of the defects observed in *ΔSslac2* mutants suggest a failure to respond to environmental factors, but it’s unknown if this failure is specific to features such as pH and surface recognition or more generalized. To address this question, an RNA sequencing analysis was conducted to compare WT (1980) and *ΔSslac2-1* strains in both defined glucose minimal media (GMM) and GMM with the addition of soybean green stem extract (GrSt) to simulate the presence of plant material. A principal component analysis was conducted to examine the variation in gene expression profiles, and unsurprisingly, a large transcriptional shift was observed in the WT strain with the addition of GrSt, reflecting the pathogen’s response to additional, proteins, carbohydrates, and plant chemical signals (Fig. 7A). In contrast, the *ΔSslac2-1* is largely non-responsive to the addition of GrSt (Fig. 7A). This is further visualized by the quantification of differentially expressed genes in *ΔSslac2-1* and WT strains (Fig. 7B, Supp. File 1).

**Figure 7.**
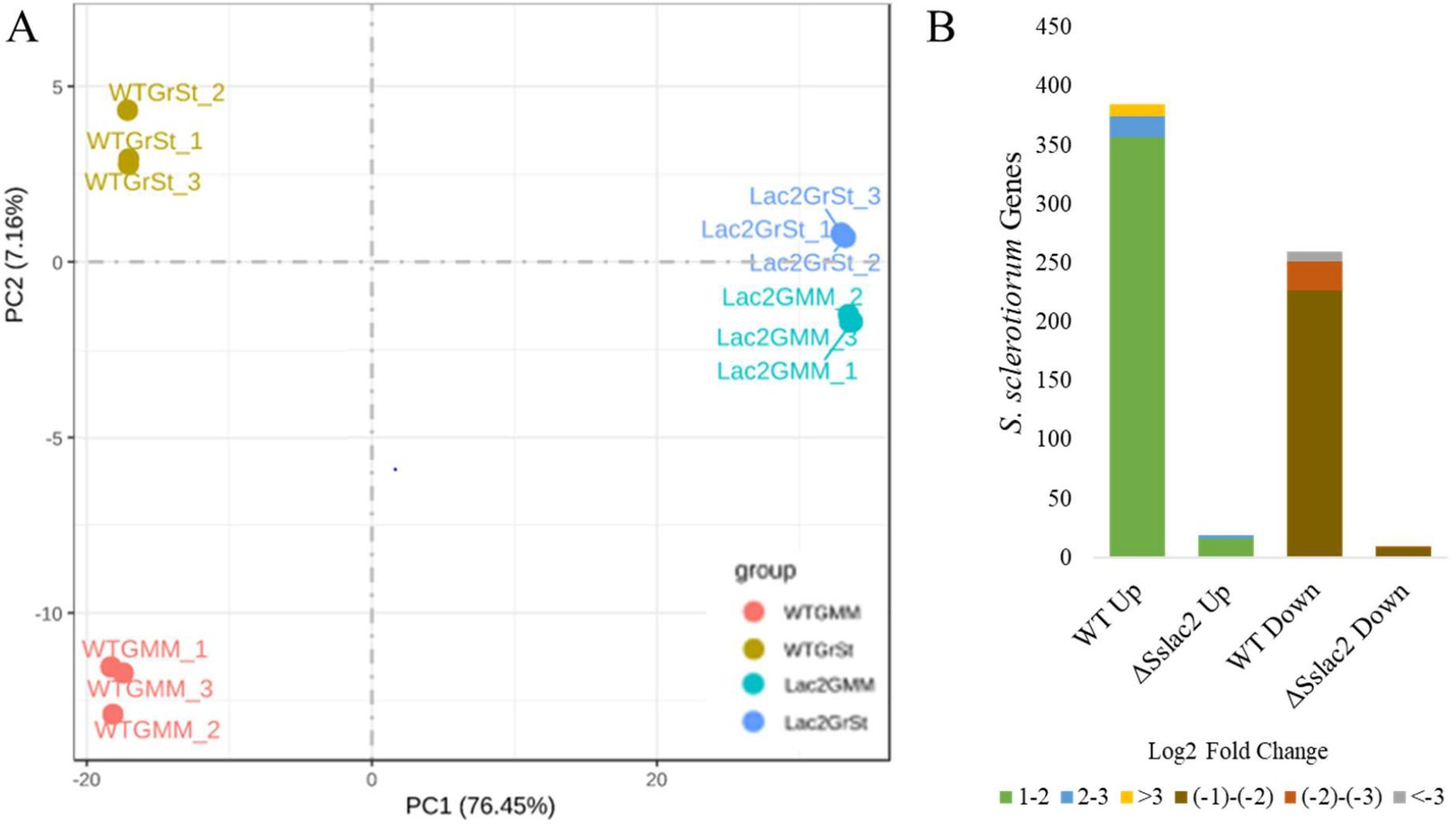
Gene expression profiles of WT (1980) and *ΔSslac2-1*. A) PCA plot comparing expression profiles of WT (1980) and *ΔSslac2-1* grown in GMM or GMM + GrSt. B) Number of genes found to be differentially regulated (FDR < 0.05; Log_2_ Fold-Change >1 or <-1) between GMM and GMM + GrSt for each strain. Up = upregulated in GMM + GrSt. Down = downregulated in GMM + GrSt.

### *ΔSslac2* environmental sensing defect is likely due to alterations in fungal cell wall structure

Laccases within ascomycete fungi have primarily been associated with detoxification and cell wall modification, so the sensitivity of *Sslac2* to antifungal plant defense compounds and cell wall stressors was assessed. Both WT (1980) and *ΔSslac2* mutants were grown on PDA plates supplemented with benzoic acid, ferulic acid, cinnamic acid, or resveratrol (Fig. 8A). The first three compounds are components of the plant phenylpropanoid pathway and are known to be toxic to *S. sclerotiorum*, whereas the latter is a well characterized phytoalexin from grapevine and known substrate of the *B. cinerea* laccase Bclcc2 (Ranjan et al. 2019; Schouten et al. 2002). *ΔSslac2* mutants were significantly more susceptible to all tested compounds. This result is somewhat surprising because the *ΔBclcc2* strain is known to show greater resistance to the phytoalexin resveratrol than WT *B. cinerea* (Fig. 8A)

**Figure 8.**
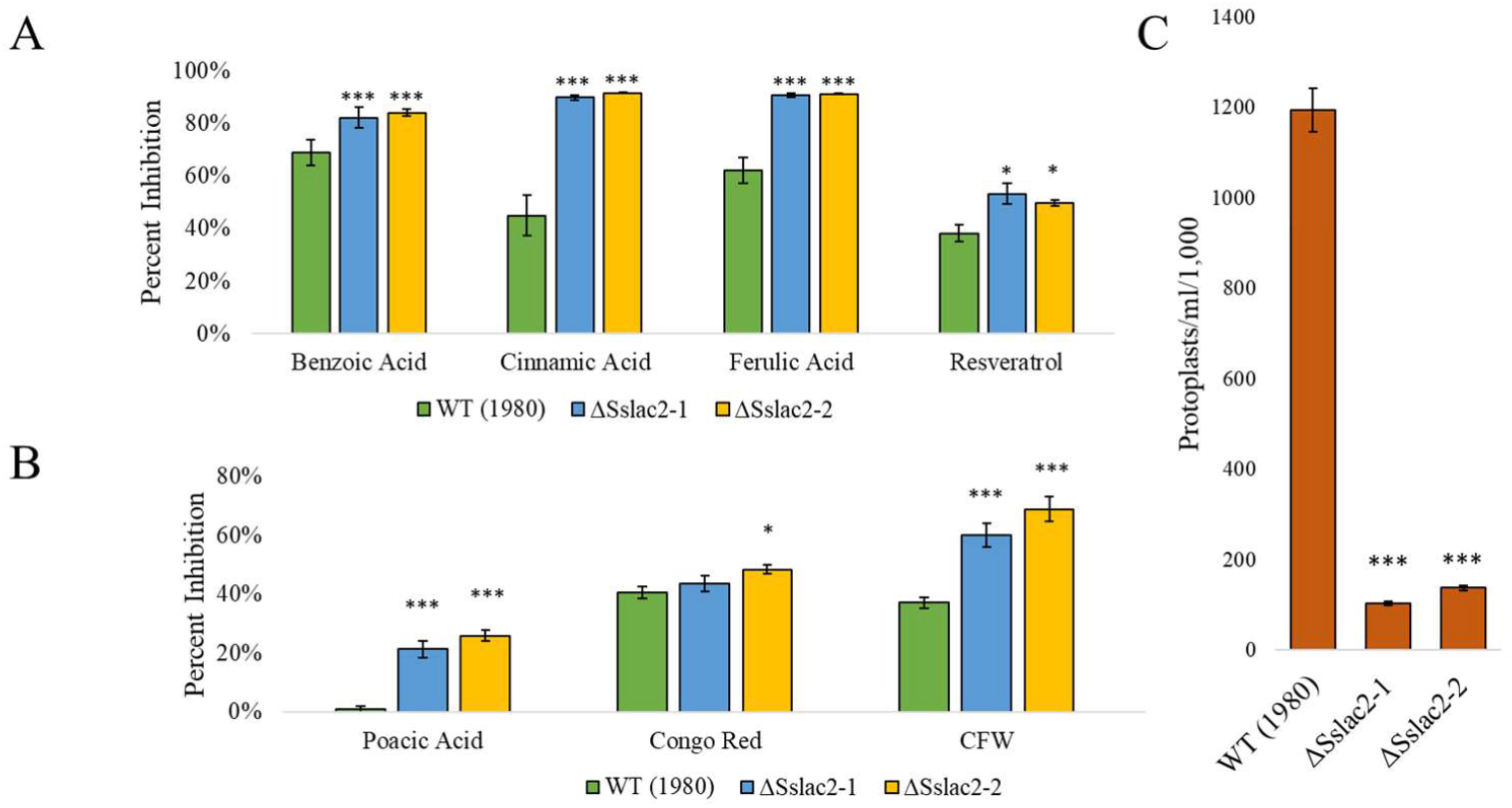
Response of the WT and mutants to chemical stresses and cell wall degrading enzymes. A) Relative inhibition of the WT and mutants when grown on plant derived antifungal compounds. Strains were grown on PDA amended with benzoic acid (150 ug/ml), cinnamic acid (150 ug/ml), ferulic acid (500 ug/ml), or resveratrol (200 ug/ml) and compared to growth on a PDA plate amended with DMSO as a control. B) Relative inhibition of the WT and mutants when grown in the presence of cell wall stressors. Strains were grown on PDA amended with poacic acid (50 ug/ml), congo red (150 ug/ml), or calcofluor white (250 ug/ml) and compared to growth on a PDA plate amended with DMSO as a control. C) Protoplast counts from WT and mutants after 3-hour incubation in lysing enzymes from *Trichoderma harzianum*. Statistical analysis utilized a Student’s t-test on three biological replicates of each strain (*<0.05, **<0.01, ***<0.001).

To assess the susceptibility of *ΔSslac2* mutants to cell wall stressors, all strains were additionally grown on plates amended with poacic acid, calcofluor white (CFW), or Congo red (Fig. 8B)(Piotrowski et al. 2015). Both mutants demonstrated significantly greater susceptibility to poacic acid and CFW and moderately increased susceptibility to Congo red (Fig. 8B). As this difference in susceptibility could be due to either alterations in hyphal cell wall architecture or a general difficulty in responding to chemical stresses, as is suggested by Fig. 8A, we attempted to form protoplasts of both strains using a cocktail of cell wall degrading enzymes from *Trichoderma harzianum*. While protoplasts could be efficiently generated from WT (1980) hyphae, *ΔSslac2* hyphae remained largely intact, suggesting that alterations to the hyphal cell wall may be reducing the efficiency of the enzyme cocktail (Fig. 8C).

Additional evidence for alterations to cell wall structure can be seen in the assessment of surface hydrophobicity through hyphal wetting. Two-week-old cultures of WT and *ΔSslac2* mutants were treated with droplets of either water or water + 0.01% Triton X-100. While both treatments were maintained as droplets on the surface of the WT mycelia, water droplets with and without Triton X-100 immediately soaked into the mutant mycelia, suggesting a dramatic decrease in hyphal hydrophobicity (Fig. S4). This alteration may help to additionally explain the accelerated growth of the mutants in liquid culture, as nutrient exchange through liquid media may be more efficient through a more hydrophilic hyphal cell wall, as was noted in hydrophobin mutants of *Trichoderma* spp. (Cai et al. 2020). An assessment of the hyphal surface was made with scanning electron microscopy, and while clear differences in the cell surface were difficult to discern, obvious changes to hyphae and hyphal growth patterns were observed (Fig. 9). While WT growth typically consists of thin hyphae branching to evenly disperse across an environment, *ΔSslac2* mutant hyphae were noticeably thicker in diameter and often found to grow in bundles (Fig. 9). This may be due to increasingly hydrophilic hypha adhering to one another and may be either a cause or an effect of the previously noted defect in growth directionality.

**Figure 9.**
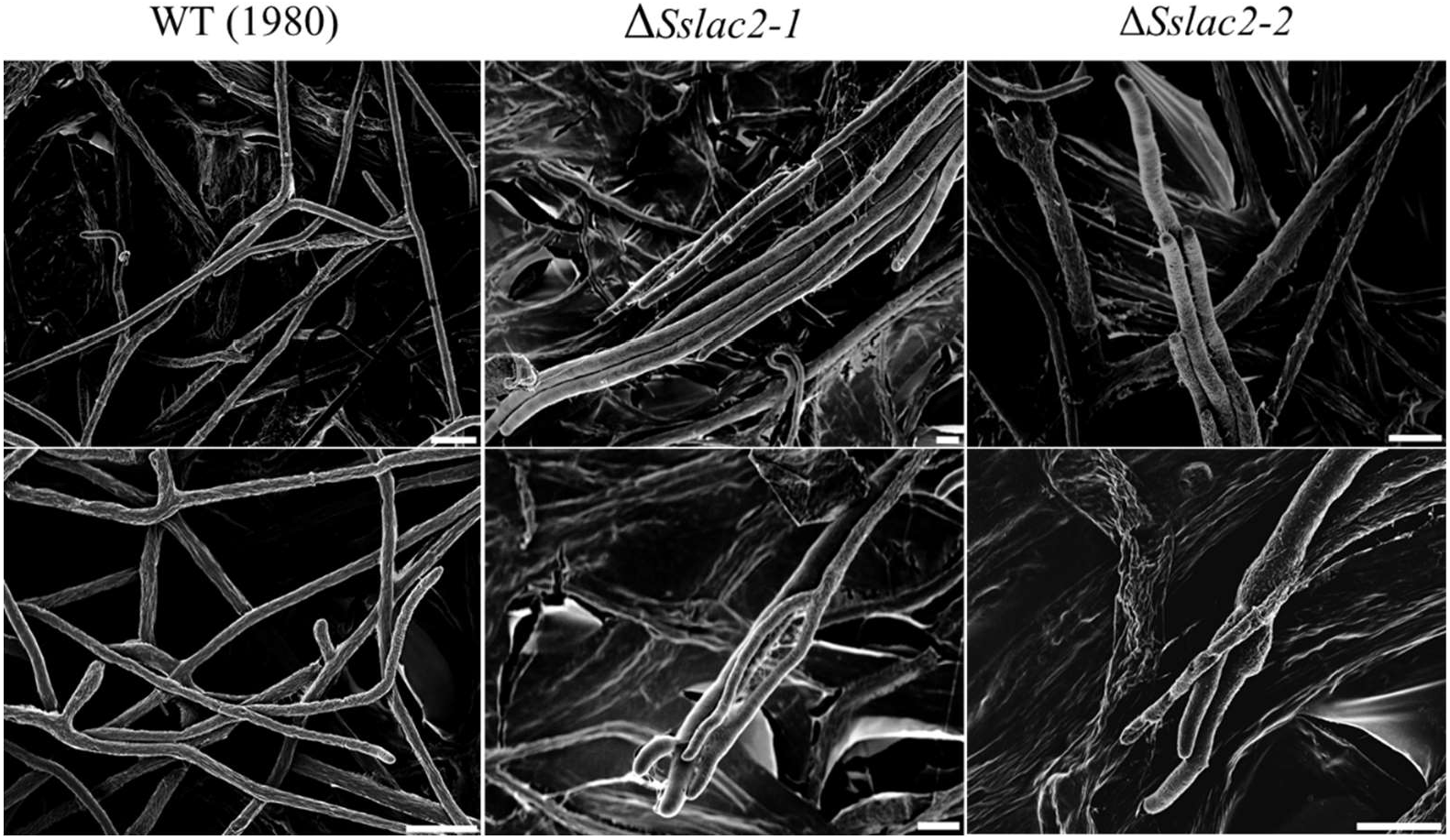
Scanning electron micrographs of WT and mutant strains of *S. sclerotiorum*. White scale bars correspond to 10 microns.

A higher resolution analysis of cell well structure was conducted through transmission electron microscopy (TEM) to assess potential changes in cell wall architecture in the mutant strains. While no structural modifications could be confirmed in the *ΔSslac2* strains given the inherent variability in *S. sclerotiorum* cell walls relative to hyphal age, a clear textural difference in the extracellular matrix coating the cells was seen (Fig. 10; Fig. S5). This apparent failure by the laccase mutants to properly assemble their exterior cell wall ultrastructure may contribute to the previously noted changes in their physiochemical properties and altered responses to external environmental signals (Fig. 8; Fig. S4).

**Figure 10.**
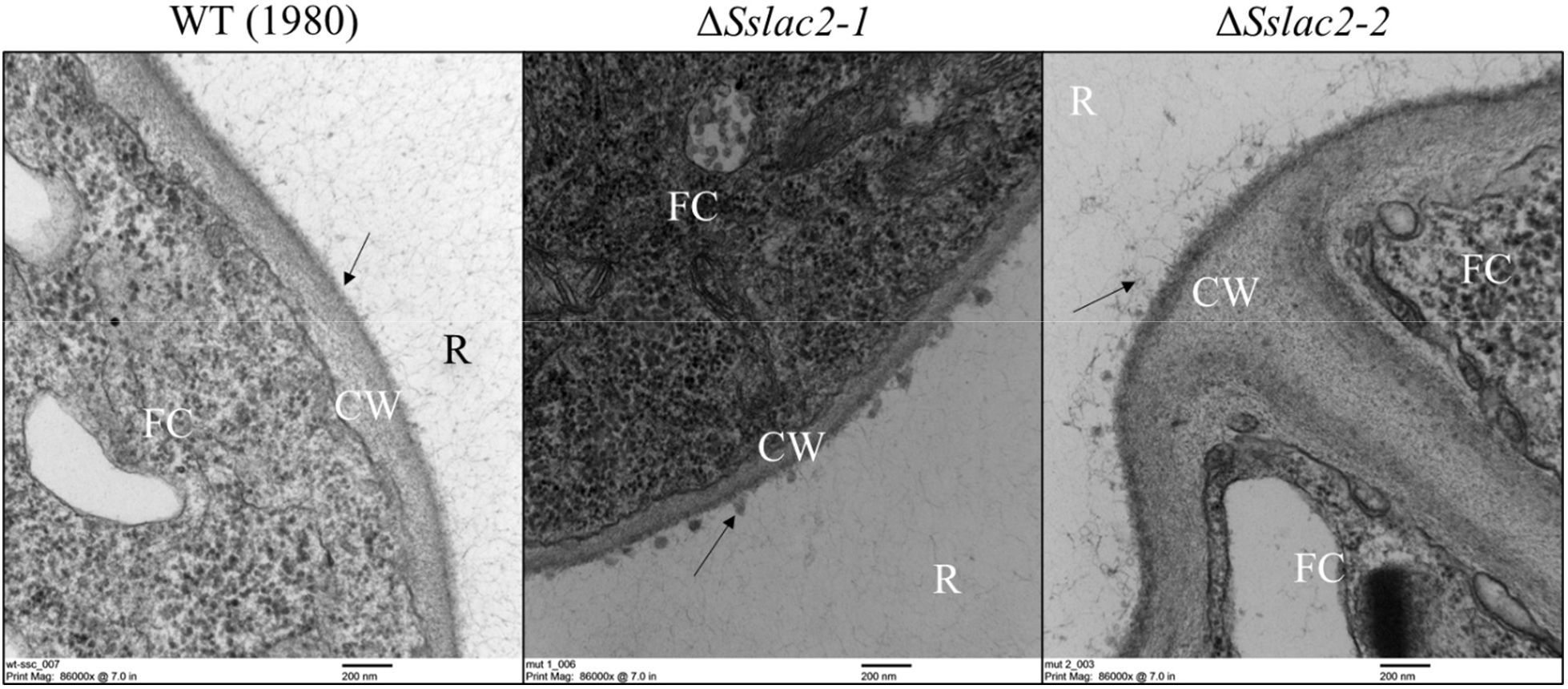
Transmission electron microscopy (TEM) photos of the cell walls and extracellular matrices of WT (1980), *ΔSslac2-1*, and *ΔSslac2-2*. R – Resin, FC – Fungal Cell, CW – Cell Wall, arrows denote the fungal extracellular matrix.

### Host-induced gene silencing (HIGS) targeting *Sslac2* induces resistance to *S*. *sclerotiorum* in soybean

Given the clear importance that *Sslac2* plays in both the virulence and development of *S. sclerotiorum*, we considered the value that this gene may have as a target of HIGS-mediated disease control. Targeted gene silencing using stable transgenic lines has been shown to provide robust disease control against *S. sclerotinia* in some pathosystems, so *Sslac2* gene silencing was first assessed using a viral vector. To achieve this, a segment of *Sslac2* was cloned into a modified Bean pod mottle virus (BPMV) vector which was then used to biolistically inoculate soybean seedlings (Zhang et al. 2010). Infected material was used to inoculate new seedlings with either empty vector (EV) or *Sslac2*-targetting BPMV, which were subsequently infected with WT *S. sclerotiorum*. Stem lesions were measured over a week of infection, and significantly smaller lesions were observed on plants in which *Sslac2* was being silenced (Fig. 11). The reddening seen on BPMV-*Sslac2* stems in response to infection indicates a successfully induced resistance response by soybeans against *S. sclerotiorum* invasion. This finding indicates that the gene silencing of *Sslac2* likely serves to limit the virulence of the pathogen, while allowing the host time to mount a more successful defense (Fig. 11A)(Ranjan et al. 2019).

**Figure 11.**
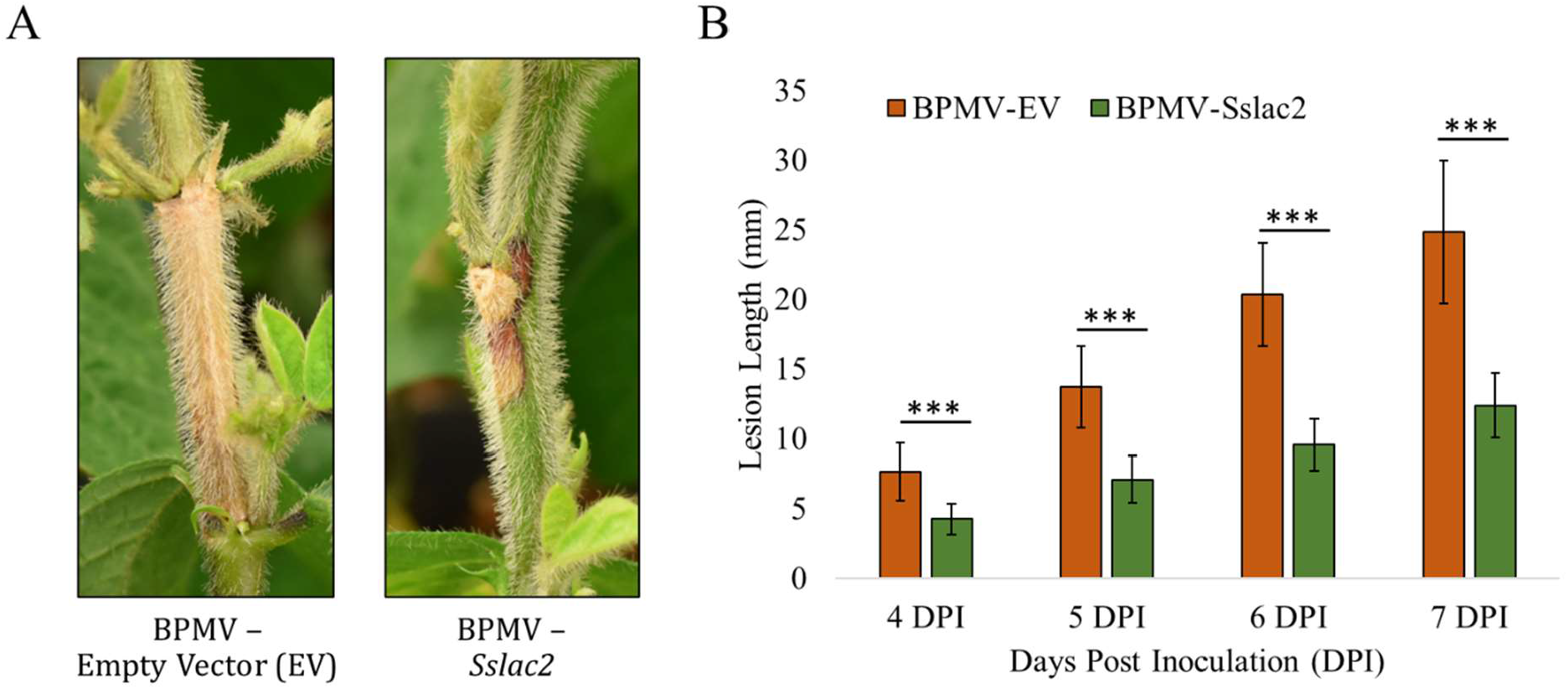
Assessment of *S. sclerotiorum* virulence in plants utilizing virus induced gene silencing (VIGS) to silence expression if *Sslac2* during infection. A) Visual appearance of lesions from *S. sclerotiorum* infection soybeans containing an empty vector (EV) strain of Bean pod mottle virus (BPMV) and soybeans containing BPMV targeting *Sslac2*. B) Quantification of lesion lengths from infection of BPMV-EV and BPMV-*Sslac2* soybeans. Statistical analysis utilized a Student’s t-test on six biological replicates of each construct. Experiment was repeated twice (*<0.05, **<0.01, ***<0.001).

## Discussion

In this study a novel laccase, *Sslac2*, was identified within the broad host-range fungal pathogen *S. sclerotiorum* and found to be critical for the proper regulation of an array of developmental and virulence traits. The defects observed from *ΔSslac2* mutants appear surprisingly expansive, but several of these defects have been noted to a lesser extent in laccase mutants of other ascomycetes (Figure 12). The mutants most closely resembling *ΔSslac2* are that of *ΔStLAC2*, from the northern corn leaf blight pathogen *S. turcica*, and *Lac1*, from the anthracnose pathogen *C. gloeosporioides*. In both cases, laccase knockout mutants were similar to *ΔSslac2* in that they had severe defects in appressorial production, leading to a loss of pathogenicity on intact plant tissue and drastically reduced virulence on wounded tissue (Ma et al. 2017; Wei et al. 2017). *ΔStLAC2* additionally displays a similar increased susceptibility to cell wall stressors and altered hydrophobicity, with an identical hyphal wetting phenotype (Ma et al. 2017). A primary difficulty in comparing the spectrum of biological roles mediated by individual laccases is that many of these functions have not been assessed across all species. In the mulberry pathogen *S. shiraiana*, knockdown mutants of the laccase *Sh-lac* generated significantly less oxalic acid than WT strains, similar to *ΔSslac2* (Lu 2017). Unfortunately, environmental acidification has not been assessed outside of these two systems, so it is unknown whether this is a common feature of laccase mutants. Nearly all of the assessed laccase mutants have some defect in melanin/pigment formation, and it is typically suggested that subsequent phenotypic alterations are due to melanogenic defects, but we argue that this explanation is unlikely in *S. sclerotiorum* (Fig. 12). *S. sclerotiorum* is known to generate melanin through the DHN melanin pathway and multiple components of this pathway have been deleted and characterized (Li et al. 2018; Y. Liang et al. 2018). Although a loss of melanization was seen in sclerotia and compound appressoria, there was no effect on fungal virulence. This agrees with expansive studies on the melanogenic genes from the closely related fungus *B. cinerea*, none of which was found to play a clear role in infection (Schumacher 2016).

**Figure 12.**
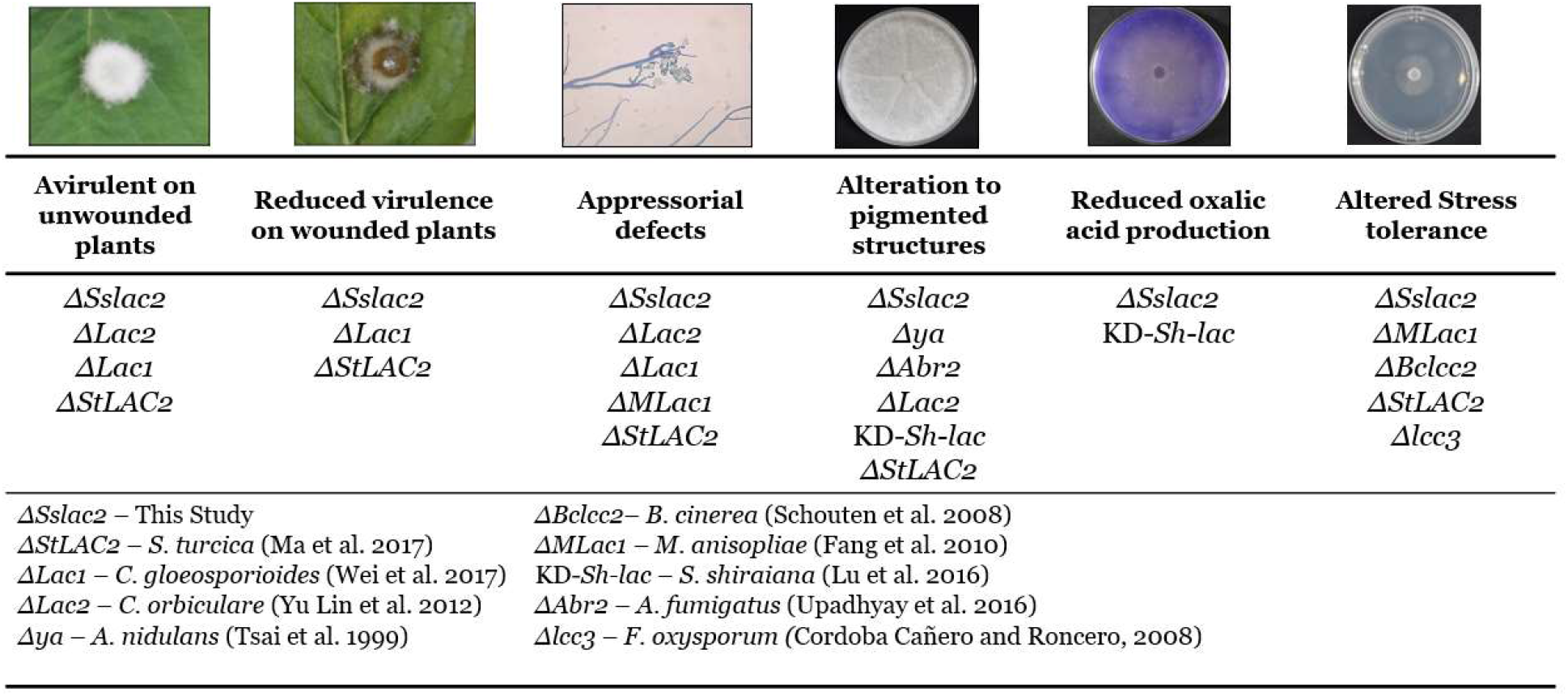
Comparison of laccase mutant phenotypes from filamentous ascomycetes.

*S. sclerotiorum* has seven putative laccases in its genome, all of which have predicted secretion signal peptides on their N-termini, but only five of which (*Sslac2-6*) contain the canonical C-terminal motif DSGx (Figure 1A; Fig. S2)(Andberg et al. 2009). While *Sslac2* is the only of these laccases with clear induction *in-planta* from our transcriptomic analysis, existing *S. sclerotiorum* expressed sequence tag libraries suggest that other laccases are expressed at distinct developmental stages (sclerotial development, carpogenic germination apothecial formation), possibly playing a more classic role in melanin deposition (Figure 1B)(Lyu et al. 2015). Given the overlapping substrate ranges of many laccases, some amount of redundancy is expected and has been demonstrated in other systems (Baldrian 2006; Saitoh, Izumitsu, Morita, Shimizu, et al. 2010). Additionally, it was demonstrated in *C. orbiculare* that *ΔLac2* mutants could be functionally complemented with orthologous laccases from related species. Subsequently, the drastic phenotype observed in *ΔSslac2* mutants is somewhat surprising, because the closest characterized ortholog of *Sslac2* is the *B. cinerea* laccase *Bclcc2*. Knockout mutants of *Bclcc2* displayed no alterations in virulence and appear to be primarily involved in oxidation of environmental phenolics, although both *ΔSslac2* and *ΔBclcc2* strains show abolished tannic acid oxidation activity (Schouten et al. 2002, 2008). Laccases are a class of multicopper oxidase requiring a core of copper to catalyze oxidation, and while no phenotypes similar to *ΔSslac2* have been observed in *B. cinerea* laccase mutants, a nearly identical phenotype has been observed in mutants of the copper transporter *BcCCC2* (Saitoh, Izumitsu, Morita, and Tanaka 2010). Knockouts of *BcCCC2* show reduced melanization, malformed and reduced compound appressoria, no pathogenicity on unwounded tissue, and reduced virulence on wounded tissue, all of which we observed in *ΔSslac2* strains. These deficiencies were attributed to the failure of the *ΔBcccc2* mutant to provide copper to copper-containing enzymes, of which laccases were major likely recipients, and suggests that *B. cinerea* may utilize laccases other than *BcLcc2* to perform similar functions to *Sslac2* (Saitoh, Izumitsu, Morita, and Tanaka 2010).

Our data show that *ΔSslac2* mutants are reduced in hydrophobicity relative to WT strains, as observed through enhanced hyphal wetting (Fig. S4). Typically, surface hydrophobicity is largely mediated by the presence of surface hydrophobins, the presence of which are both ubiquitous among and unique to filamentous fungi (Linder et al. 2005). It’s surprising that surface hydrophobicity would be affected after the deletion of a single laccase because the *S. sclerotiorum* hydrophobins should still be intact; however, TEM analysis of the WT and *ΔSslac2* strains suggests that the extracellular matrix (ECM) of mutant strains may be more intrinsically disordered than the WT (Fig. 10; Fig. S5). Such a change in the ECM, the component of hyphae that interacts most directly with the environment, may help to explain why the mutant hyphae were observed to “stick” to one another and why hydrophobins may by incapable of maintaining hydrophobicity. This loss of hydrophobicity may also explain the enhanced growth of the mutants in liquid culture, as a similar phenotype has been observed in more hydrophilic strains of *Trichoderma* (Cai et al. 2020). Additionally, modifications to the ECM could undermine fungal cell receptor activity, which could help to explain the aberrant environmental sensing phenotypes observed in this study. This defect in environmental sensing might also explain many of the developmental phenotypes we observed, including the observed reduction in oxalic acid secretion and canonical compound appressoria formation (Fig. 5). This study is the first to provide evidence for roles of fungal laccases in ECM formation and environmental sensing.

Other Sslac2-driven modifications to cell wall composition are likely, as we show that *ΔSslac2* is very resistant to protoplasting by a cocktail of cell wall degrading enzymes (Fig. 8C). Such a phenotype was additionally observed in knockout mutants of *Rho1*, a GTPase, from *Fusarium oxysporum*, which displayed strikingly similar developmental defects to *ΔSslac2*, including a growth defect specific to solid surfaces, a loss of pathogenicity on plants, increased susceptibility to cell wall stressors, and resistance to protoplasting (Martínez-Rocha et al. 2008). Broadly, *Rho1* orthologues in yeast are known to play a role in polarized cell growth through regulation of the actin cytoskeleton and can directly interact with the β-1,3-glucan synthase in fungal cell walls (Martínez-Rocha et al. 2008). This activity likely extends to filamentous fungi as well, given the growth morphology and cell wall alterations observed in *ΔRho1* from *F. oxysporum*) and *ΔRhoA* (*A. nidulans*), and supports a connection between cell wall composition and the thigmotropic phenotypes observed in *ΔSslac2* (Fig. 7,9; Fig. S3-5)(Guest, Lin, and Momany 2004; Martínez-Rocha et al. 2008). The precise interplay between cell wall biosynthesis and thigmotropism is currently unclear, but such alterations may additionally play a role in the avirulent phenotype of *ΔSslac2* strains, as plants are known to activate defenses in response to fungal cell wall components (Fig. 6)(Pieterse et al. 2012). If the cell wall of *ΔSslac2* mutants were modified in a way that increases the release of such components, as was predicted in *ΔRho1*, then a reduction in virulence would be expected.

Given the clear and pivotal role that *Sslac2* plays in pathogenesis, we chose to evaluate it as a target of host induced gene silencing (HIGS) to achieve disease resistance in soybeans. An initial screen using a viral vector confirmed that silencing *Sslac2* significantly increases plant resistance to *S. sclerotiorum* (Fig. 11). A drawback of such an approach is that viral vectors are often only capable of partial gene silencing, given the relatively low viral titer in plants used for this assay (Ranjan et al. 2019). In the future, stable transgenic soybean lines expressing hairpin dsRNA targeting *Sslac2* will be generated and evaluated for resistance to *S. sclerotiorum* infection.

In summary, this study characterizes a fungal laccase critical for proper development, virulence, and environmental sensing in the broad host-range fungal plant pathogen *S. sclerotiorum*. Future work will focus on elucidating the chemical substrates of *Sslac2* and precise mechanisms by which this protein mediates fungal thigmotropism and responses to environmental stimuli. Efforts will also focus on evaluating *Sslac2* and other fungal laccases as targets of gene silencing for disease control.

## Materials and Methods

### Plant and Fungal Growth

All soybean and *Nicotiana benthamiana* plants were maintained in the greenhouse or growth chamber at 24 ± 2 °C with 16-h light/8-h dark photoperiod cycle. Plants were watered daily and supplemented with fertilizer (Miracle-Gro) every week.

All *S. sclerotiorum* cultures were maintained on potato dextrose agar (PDA) plates or PDA supplemented with 50 ug/ml hygromycin in the case of knockout strains. Liquid cultures were grown in potato dextrose broth. Cellophane assays were conducted by autoclaving pre-cut rings of cellophane prior to being placed on 100×15mm petri dishes containing PDA.

### Fungal Transformation

Gene knockouts were generated in *S. sclerotiorum* using a CRISPR-Cas9 method in combination with a modified form of the Rollins et al., 2003 protocol. Split-markers targeting *Sslac2* were generated using polymerase chain Reaction (PCR) by amplifying 5-600 bp regions upstream (Sslac2-LF-F and Sslac2-LF-R) and downstream (Sslac2-RF-F and Sslac2-RF-R) regions of *Sslac2*. These amplicons were designed to contain 20 bp sequences with homology to the 5’ and 3’ regions, respectively, of the hygromycin resistance cassette (HygR; 1.8 kb) found in pCRISPR-Cas9-TrpC-Hyg(Li et al. 2018). The hygromycin resistance cassette was amplified using The two flanking regions and the HygR were connected through fusion PCR as described in Szewczyk et al., 2007 for a product of ∼3kb (Szewczyk et al. 2006). Split markers were generated from this product by using primers internal to HygR (Hyg Split F and Hyg Split R) in conjunction with Sslac2-RF-F and Sslac2-RF-R, yielding two amplicons with an overlapping region of ∼400 bp.

Two small guide RNAs (sgRNAs) targeting *Sslac2* were designed using the E-CRISP Design Tool (http://www.e-crisp.org/E-CRISP/index.html), generated using the GenCrispr sgRNA Screening Kit (L00689; Genscript Biotech Corp.), and diluted to a concentration of 4 uM. Alt-R S.p. Cas9 nuclease 3NLS (1081058; IDT) was diluted to a concentration of 4 uM and combined with sgRNA at a 1.2-to-1 ratio (3.6 ul of sgRNA to 3 ul of Cas9 protein) and incubated at room temperature for 5 minutes to assemble the RNP complex. These complexes were combined with 1 ug of each split-marker and transfected into *S. sclerotiorum* protoplasts using the polyethylene glycol (PEG) transformation described in Rollins et al., 2003(Rollins 2003).

Transformants capable of surviving on PDA containing 50 ug/ml hygromycin were subjected to 5 rounds of hyphal tipping before undergoing DNA extraction to confirm the replacement of Sslac2 with the HygR marker. Primers internal and external to Sslac2 and HygR were used to confirm deletion and primers targeting *Histone 3* (H3 F and H3 R) were used as a control (Fig S6: Table S1). DNA was extracted using the cetyl trimethyl ammonium bromide (CTAB) method described in Talbot et al, 1993.

### Virus-induced gene silencing (VIGS) assay and construct generation

A modified Bean pod mottle virus (BPMV) vector was used to assess *Sslac2* as a potential target of VIGS for disease control(Zhang et al. 2010). In order to silence *Sslac2* (Sscle_03g023030; SS1G_00974; XM_001598835), a 267 base pair sequence was selected within the mRNA of S. sclerotiorum strain 1980 (GenBank Accession XM_001590428). Total RNA was extracted from *S. sclerotiorum* using the Maxwell® RSC Plant RNA Kit, and cDNA was generated using an AMV first strand cDNA synthesis kit (New England Biolabs, Catalog # E6550). The segment was amplified through PCR with PstI and BamHI restriction sites incorporated onto the double-stranded cDNA using specific primers (PstI-VIGS-F and BamHI-VIGS-R) (Supplementary Table 1). The amplicon underwent gel purification (QIAquick Gel Extraction Kit®, QIAGEN), then both the amplicon and the viral vector RNA2 plasmid (pBPMV-IA-V1) were subjected to restriction digestion with PstI/BamHI before being ligated together to form BPMV-Sslac2(Zhang et al. 2010). The vector plasmids were then transformed into DH5α competent cells using 5 μl of the purified ligation product per 50 μl of competent cells, a 30 min ice incubation, 45 s heat shock in a 42°C water bath, and incubation in 500 μl of Luria broth (LB) for 1 h at 37°C. Using glycerol stocks, midi preparations were conducted (Fast Ion Plasmid Midi Kit®, IBI Scientific) for subsequent biolistic inoculations. Biolistic inoculations were performed as described in McCaghey and Shao et al, 2021 (McCaghey et al. 2021).

### Plant Disease Assays

Soybeans: For detached leaf assays, leaves were taken from the first trifoliate of 5–6-week-old plants (cv. Williams 82) and placed in petri dishes containing two layers of filter paper/paper towel and 7 milliliters of sterile water. Leaves were inoculated near the center with agar plugs of actively growing wild-type (1980) or mutant *S. sclerotiorum*. Damaged leaves were scored four times in a crisscross pattern using a sterile scalpel directly under the agar plug. Petri dishes were wrapped in parafilm, placed at room temperature, and photographed every twenty-four hours. All samples were inoculated in triplicate.

For VIGS assays, 10-14 day old plants (cv. Traff) were rub-inoculated with lyophilized leaves infected with BPMV-Sslac2 as described in McCaghey and Shao et al, 2021 (McCaghey et al. 2021). Plants were then allowed to grow an additional 5 weeks prior to being inoculated with WT (1980) *S. sclerotiorum* through cut petioles. Briefly, deep-well plates (100mm x 55mm) containing 75 ml of PDA were inoculated with WT (1980) and allowed to grow for two days. The petiole of the first trifoliate was cut ∼2 cm from the main stem with a razor and a plug from the leading edge of mycelia was punctured using an inverted one-ml pipette tip. The inverted pipette tip with agar plug, was then slid onto the excised petiole. Lesions were measured with digital calipers 4–7 Days post-inoculation (DPI). Three plants were inoculated per 1 L pot and 5 pots were tested for both the empty vector and BPMV-Sslac2 infected plants. The statistical significance of lesion differences was assessed with a Student’s t-test (*<0.05, **<0.01, ***<0.001). The experiment was replicated twice.

*N. benthamiana*: Leaves were taken from 6-7 week old plants and placed in petri dishes containing two layers of filter paper/paper towel and 7 milliliters of sterile water. Leaves were inoculated near the center with agar plugs of actively growing wild-type (1980) or mutant *S. sclerotiorum*. Damaged leaves were scored four times in a crisscross pattern using a sterile scalpel directly under the agar plug. Petri dishes were wrapped in parafilm, placed at room temperature, and photographed every twenty-four hours.

### Stress Testing

All stress test assays were performed in 60 mm x 15 mm petri dishes containing PDA supplemented with benzoic acid (150 ug/ml), cinnamic acid (150 ug/ml), ferulic acid (500 ug/ml), resveratrol (200 ug/ml), congo red (50 ug/ml), calcofluor white (250 ug/ml), or poacic acid (50 ug/ml). Cultures were allowed to grow for 24-48 hours prior to being photographed. Colony areas were quantified in ImageJ(Schneider, Rasband, and Eliceiri 2012). All samples were tested in triplicate and the statistical significance of colony area differences was assessed with a Student’s t-test (*<0.05, **<0.01, ***<0.001).

### Protoplasting Assay

Protoplasting was done using a modified protocol from Rollins et al, 2003. Briefly, three agar plugs of each strain were grown for two days in petri dishes containing PDB at room temperature. Agar plugs were excised with tweezers and a scalpel, and each sample was washed with water and then protoplast buffer (0.8 M MgSO₄·7H₂O, 0.2 M Sodium citrate·2H₂O, pH 5.5). Samples were roughly chopped with a sterile razor blade and placed in 17 ml of protoplast buffer. For each sample, 100 mg of lysing enzyme from *Trichoderma harzianum* (Sigma Aldrich, L1412) was dissolved in 3 ml of Novozyme buffer (1 M Sorbitol, 50 mM Sodium citrate·2H₂O) and then filtered through a 0.45 um filter directly into to protoplast buffer containing the sample. All samples were incubated in a 28C shaker at 120 RPM for 3 hours before being filtered through four layers of Miracloth to collect protoplasts. Samples were centrifuged at 3000xg for 10 minutes to pellet protoplasts, which were subsequently reconstituted in 1 ml protoplast buffer before quantification with a hemocytometer. All samples were run in triplicate. The statistical significance of protoplast count differences was assessed with a Student’s t-test (*<0.05, **<0.01, ***<0.001). The experiment was replicated twice.

### Laccase activity assay

Laccase Activity was assessed using either 0.2 mM 2,2’-azino-bis(3-ethylbenzothiazoline-6-sulfonic acid (ABTS), 0.2 mM ABTS + 0.6 mM CuSO_4_, or 2.5 mg/ml tannic acid amended to PDA plates (100 mm x 15 mm)(Ma et al. 2017; Schouten et al. 2002). Agar plugs of actively growing hyphae from each strain was used to inoculate the center of plates and were allowed to grow for 24 hours prior to being photographed. Laccase activity was associated with the accumulation of bluish/purple pigments in the case of ABTS and brown pigment in the case of tannic acid. CuSO_4_ amendment was utilized as a known inducer or laccase activity(Buddhika, Savocchia, and Steel 2021).

### Hydrophobicity assay

Three-week-old cultures of all strains were topped with100 ul of either H₂O or H₂O + 0.01% Triton X-100. Photos were taken 2 minutes after treatment to observe mycelial soaking.

### Compound Appressoria Observation/Quantification

Agar plugs of actively growing mycelium were collected and placed face down on glass slides and then incubated in the dark at room temperature in a sealed container overnight (∼16 hours). A scalpel was used to cut the agar plugs away from hyphae which has grown onto the glass slide and the plugs were removed. Mycelia were stained with 0.05% trypan blue for 1 hour before being rinsed with water to remove the dye. As the mutant strains are substantially less hydrophobic than the wild type and therefore attach poorly to the glass slide, the water rinse was done by carefully removing the dye and replacing it with water. This process was repeated 10x for each agar plug. Stained mycelia were then observed under a compound scope to observe the production of compound appressoria. The statistical significance of compound appressorium count differences was assessed with a Student’s t-test (*<0.05, **<0.01, ***<0.001).

### RT-PCR of *Sslac2* on PDA and PDB

WT (1980) *S. sclerotiorum* was grown for 48 hours at room temperature either in a 125 ml Erlenmeyer flask containing PDB on a rotary shaker (120 RPM) or on a plate of PDA (100 mm x 15 mm). Samples in PDB were then removed from the broth and flash frozen in liquid nitrogen and ground into a powder with a mortar and pestle. Samples on PDA had liquid nitrogen poured directly onto plates to flash freeze mycelium and the underlying PDA, then a thin layer of mycelia was scraped off using a pre-chilled scalpel. Total RNA was extracted from frozen mycelia using the Maxwell® RSC Plant RNA Kit, and cDNA was amplified using an AMV first strand cDNA synthesis kit and was normalized to 50 ng/ul for each sample (New England Biolabs, Catalog # E6550). *Sslac2* was amplified using specific detection primers (Sslac2 Det F and R) for 30 cycle

### Scanning and Transmission Electron Microscopy (SEM and TEM)

For SEM samples were grown on PDA plates embedded with 10 mm Whatman filters shortly after plates were poured, allowing for a thin layer of agar to cover the filters. Samples were grown for 2 days until they had completely covered the filter paper, before being submerged in a chemical fixative (78% - Ultrapure ddH2O, 10% - 10x PBS, 10% - 37% formaldehyde, 2% - 50% glutaraldehyde) overnight. Samples were treated with 1% osmium tetroxide for 30 min at 22°C. Samples were subsequently washed with a series of increasing ethanol concentrations (30 to 100% [vol/vol]), followed by critical point drying and coating with platinum. Scanning electron microscopy (SEM) of samples was performed using a LEO 1530 microscope. TEM samples were grown on PDA overlayed with cellophane for two days before 10 mm circles were peeled off and placed in the above fixative overnight. Sample preparation, sectioning, and imaging was conducted by the UW Madison Medical School Electron Microscope Facility on a Philips CM120 STEM.

## Supplementary Tables

**Supplementary Table 1.**
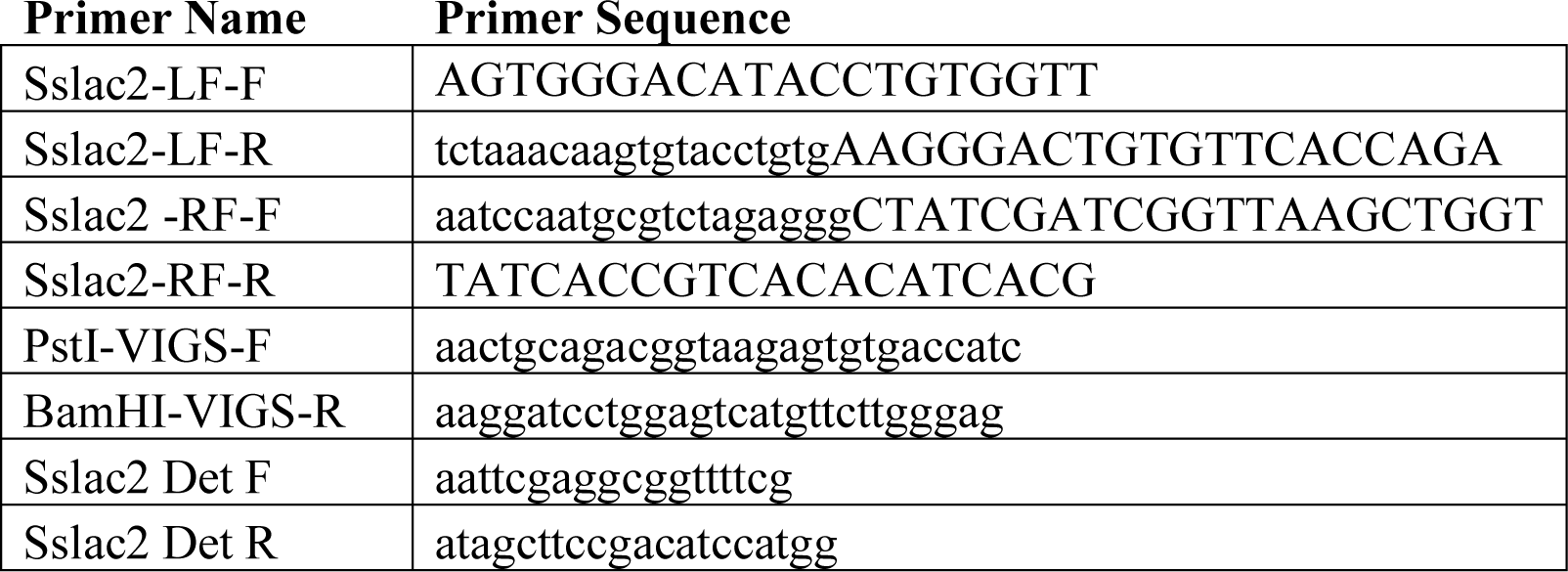
Primers used in this study.

## Supplementary Figures

**Supplementary Figure 1.**
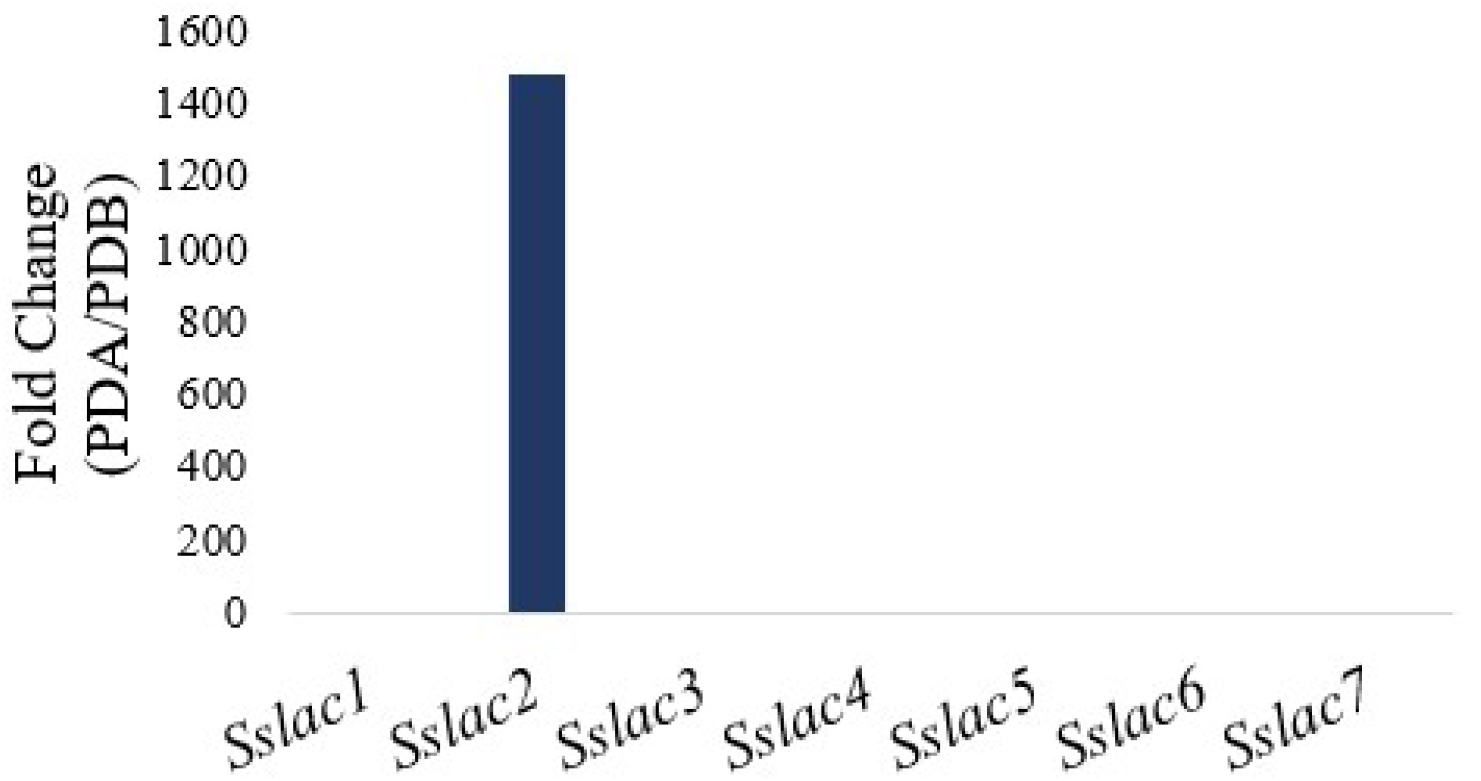
Relative transcript fold change of the seven laccases identified in the *S. sclerotiorum* genome when grown on PDA when compared to growth on PDB. Fold change values were taken taken from the transcriptomic analysis performed in Peyraud et al., 2019.

**Supplementary Figure 2.**
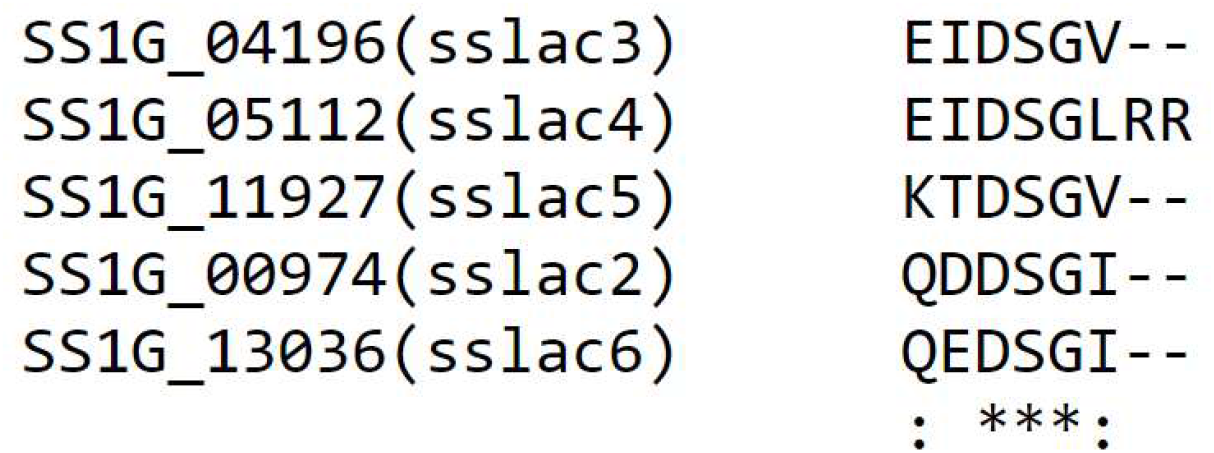
Alignment of the C-terminal amino acids from *Sslac2-6*. A conserved DSGx motif is observed at or near the C-terminus of all proteins. *Sslac1* and *Sslac7* are not included as they were missing this motif.

**Supplementary Figure 3.**
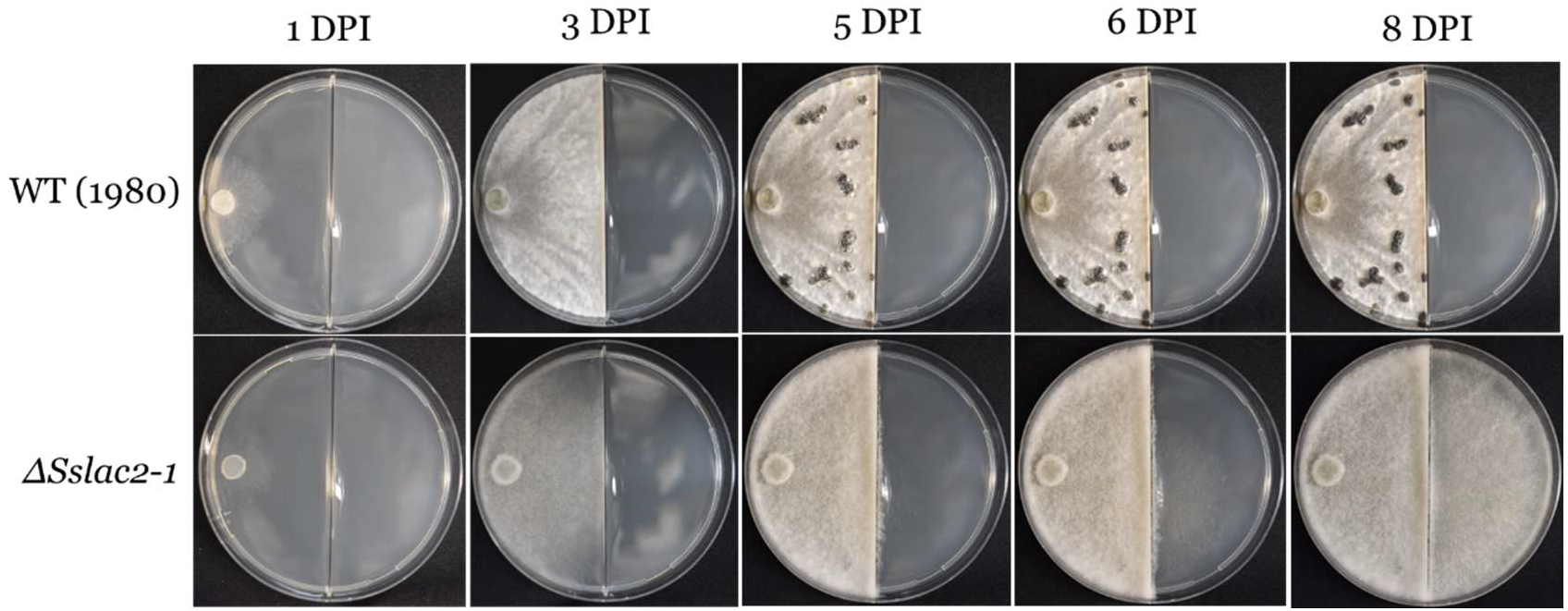
Growth of WT (1980) and *ΔSslac2-1* on a PDA split plate over 8 days post inoculation (DPI).

**Supplementary Figure 4.**
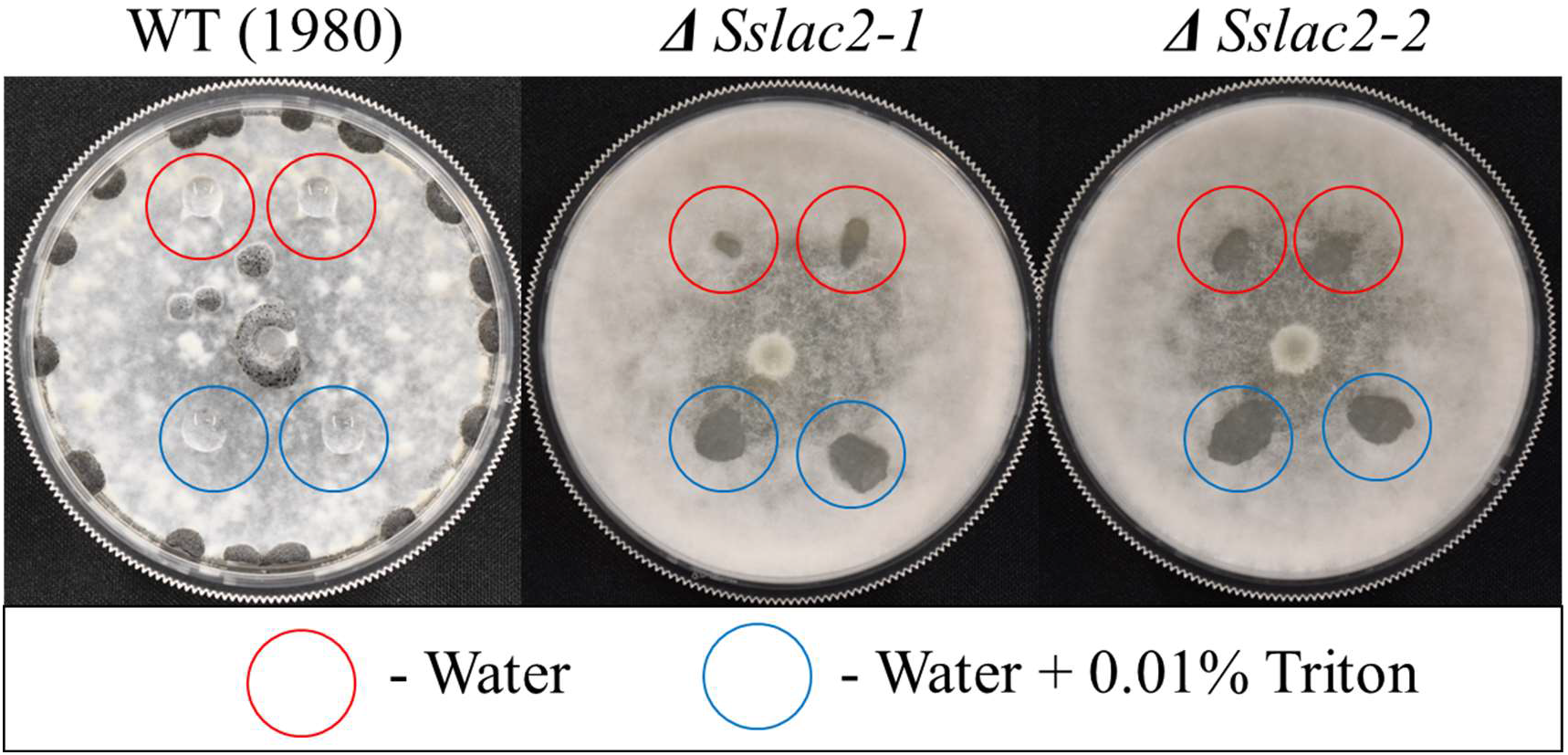
Water soaking phenotype of WT and mutant strains of *S. sclerotiorum*. 100 ul of water or water + 0.01% Triton were added to 2-week-old colonies of WT and mutant strains. Photos were taken immediately after droplets were placed. WT (1980) *!J*.*Sslac2-1 !J*.*Sslac2-2*

**Supplementary Figure 5.**
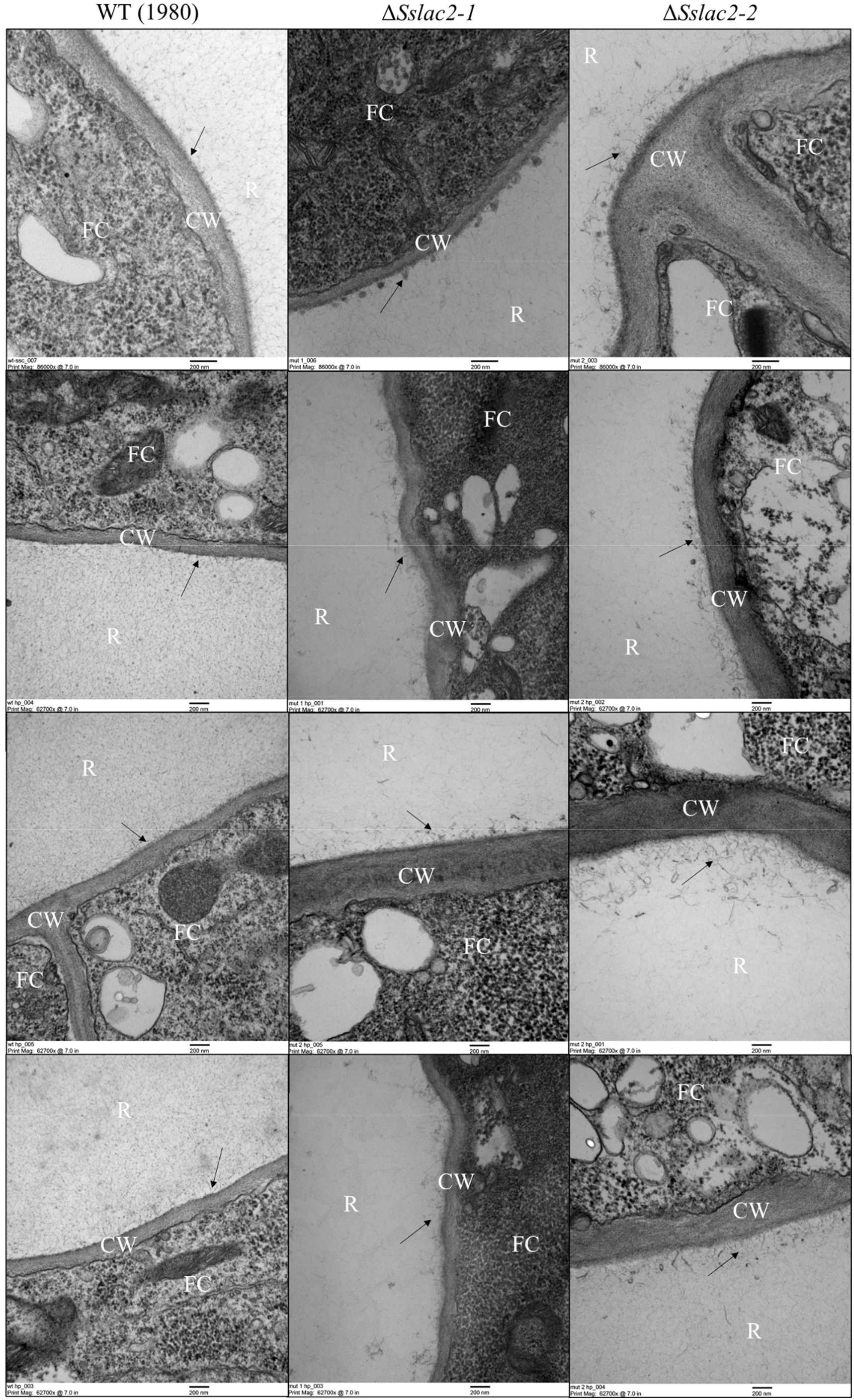
Transmission electron microscopy (TEM) photos of the cell walls and extracellular matrices of WT (1980), *ΔSslac2-1*, and *ΔSslac2-2*. R – Resin, FC – Fungal Cell, CW – Cell Wall, arrows denote the fungal extracellular matrix.

**Supplementary Figure 6.**
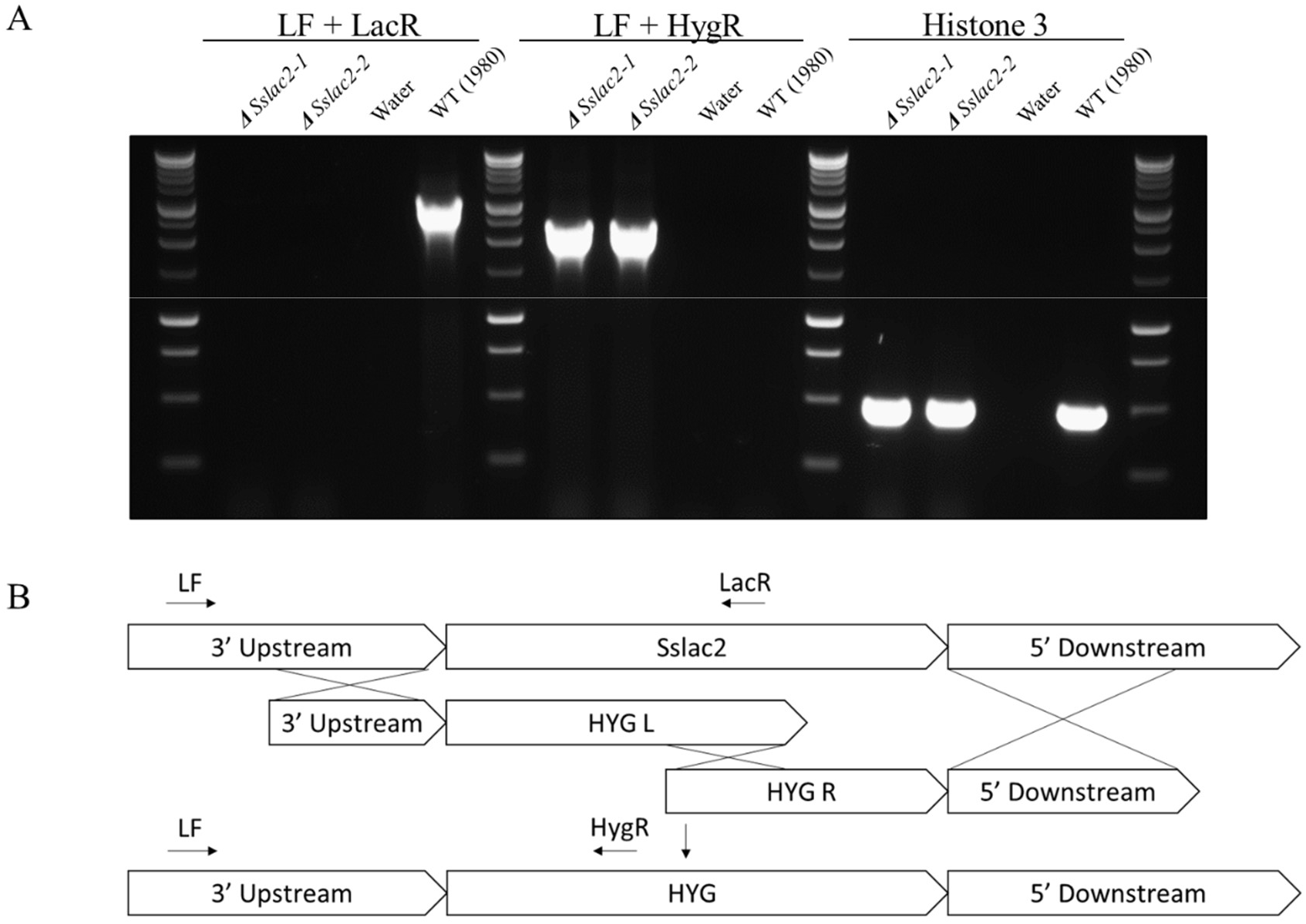
Validation of *Sslac2* gene knockouts in *S. sclerotiorum* by PCR. A) Left: PCR with external (LF) and internal (LacR) *Sslac2* primers, Middle: PCR with an external *Sslac2* primer (LF) and an internal (HygR) hygromycin resistance cassette (HYG) primer, Right: PCR of *S. sclerotiorum* histone 3 gene. B) Schematic diagram of split marker gene replacement used to generate *Sslac2* knockout strains. Approximate placement of validation primers are included.

## Citations

Adachi, Kiichi, and John E. Hamer. 1998. “Divergent CAMP Signaling Pathways Regulate Growth and Pathogenesis in the Rice Blast Fungus Magnaporthe Grisea.” Plant Cell 10(8): 1361–73.

Andberg, Martina et al. 2009. “Essential Role of the C-Terminus in Melanocarpus Albomyces Laccase for Enzyme Production, Catalytic Properties and Structure.” FEBS Journal 276(21): 6285–6300.

Arregui, Leticia et al. 2019. “Laccases: Structure, Function, and Potential Application in Water Bioremediation.” Microbial Cell Factories 18(1): 1–33. https://doi.org/10.1186/s12934-019-1248-0.

Baldrian, Petr. 2006. “Fungal Laccases-Occurrence and Properties.” FEMS Microbiology Reviews 30(2): 215–42.

Buddhika, U. V.A., S. Savocchia, and C. C. Steel. 2021. “Copper Induces Transcription of BcLCC2 Laccase Gene in Phytopathogenic Fungus, Botrytis Cinerea.” Mycology 12(1): 48– 57. https://doi.org/10.1080/21501203.2020.1725677.

Cai, Feng et al. 2020. “Evolutionary Compromises in Fungal Fitness: Hydrophobins Can Hinder the Adverse Dispersal of Conidiospores and Challenge Their Survival.” ISME Journal 14(10): 2610–24. http://dx.doi.org/10.1038/s41396-020-0709-0.

Cordoba Cañero, D., and M. I.G. Roncero. 2008. “Functional Analyses of Laccase Genes from Fusarium Oxysporum.” Phytopathology 98(5): 509–18.

Fang, Weiguo et al. 2010. “A Laccase Exclusively Expressed by Metarhizium Anisopliae during Isotropic Growth Is Involved in Pigmentation, Tolerance to Abiotic Stresses and Virulence.” Fungal Genetics and Biology 47(7): 602–7. http://dx.doi.org/10.1016/j.fgb.2010.03.011.

Feng, B Z, P Q Li, L Fu, and X M Yu. 2015. “Exploring Laccase Genes from Plant Pathogen Genomes: A Bioinformatic Approach.” Genetics and Molecular Research 14(4): 14019–36.

Guest, Gretel M., Xiaorong Lin, and Michelle Momany. 2004. “Aspergillus Nidulans RhoA Is Involved in Polar Growth, Branching, and Cell Wall Synthesis.” Fungal Genetics and Biology 41(1): 13–22.

Ten Have, Rimko, and Pauline J.M. Teunissen. 2001. “Oxidative Mechanisms Involved in Lignin Degradation by White-Rot Fungi.” Chemical Reviews 101(11): 3397–3413.

Li, Jingtao et al. 2018. “Introduction of Large Sequence Inserts by CRISPR-Cas9 to Create Pathogenicity Mutants in the Multinucleate Filamentous Pathogen Sclerotinia Sclerotiorum.” mBio 9(3): 1–19.

Liang, Xiaofei et al. 2015. “Oxaloacetate Acetylhydrolase Gene Mutants of Sclerotinia Sclerotiorum Do Not Accumulate Oxalic Acid, but Do Produce Limited Lesions on Host Plants.” Molecular Plant Pathology 16(6): 559–71.

Liang, Yue, Wei Xiong, Siegrid Steinkellner, and Jie Feng. 2018. “Deficiency of the Melanin Biosynthesis Genes SCD1 and THR1 Affects Sclerotial Development and Vegetative Growth, but Not Pathogenicity, in Sclerotinia Sclerotiorum.” Molecular Plant Pathology 19(6): 1444–53.

Lin, Shao Yu et al. 2012. “LAC2 Encoding a Secreted Laccase Is Involved in Appressorial Melanization and Conidial Pigmentation in Colletotrichum Orbiculare.” 25(12): 1552–61.

Linder, Markus B., Géza R. Szilvay, Tiina Nakari-Setälä, and Merja E. Penttilä. 2005. “Hydrophobins: The Protein-Amphiphiles of Filamentous Fungi.” FEMS Microbiology Reviews 29(5): 877–96.

Lu, Zhiyuan. 2017. “Laccase Gene Sh-Lac Is Involved in the Growth and Melanin Biosynthesis of Scleromitrula Shiraiana.” 107(3): 353–61.

Lyu, Xueliang et al. 2015. “A ‘Footprint’ of Plant Carbon Fixation Cycle Functions during the Development of a Heterotrophic Fungus.” Scientific Reports 5(July): 1–13. http://dx.doi.org/10.1038/srep12952.

Ma, Shuangxin et al. 2017. “The StLAC2 Gene Is Required for Cell Wall Integrity, DHN-Melanin Synthesis and the Pathogenicity of Setosphaeria Turcica.” Fungal Biology 121(6– 7): 589–601. http://dx.doi.org/10.1016/j.funbio.2017.04.003.

Martínez-Rocha, Ana Lilia et al. 2008. “Rho1 Has Distinct Functions in Morphogenesis, Cell Wall Biosynthesis and Virulence of Fusarium Oxysporum.” Cellular Microbiology 10(6): 1339–51.

McCaghey, Megan et al. 2021. “Host-Induced Gene Silencing of a Sclerotinia Sclerotiorum Oxaloacetate Acetylhydrolase Using Bean Pod Mottle Virus as a Vehicle Reduces Disease on Soybean.” Frontiers in Plant Science 12(July): 1–13.

Mccaghey, Megan, Jaime Willbur, Damon L Smith, and Mehdi Kabbage. 2018. “The Complexity of the Sclerotinia Sclerotiorum Pathosystem in Soybean: Virulence Factors, Resistance Mechanisms, and Their Exploitation to Control Sclerotinia Stem Rot.”

Morozova, O V, G P Shumakovich, S V Shleev, and Ya I Yaropolov. 2014. “Laccase – Mediator Systems and Their Applications: A Review.” Applied Biochemistry and Microbiology 43(5): 523–35.

Peyraud, Rémi, Malick Mbengue, Adelin Barbacci, and Sylvain Raffaele. 2019. “Intercellular Cooperation in a Fungal Plant Pathogen Facilitates Host Colonization.” Proceedings of the National Academy of Sciences of the United States of America 116(8): 3193–3201.

Pieterse, Corné M.J. et al. 2012. “Hormonal Modulation of Plant Immunity.” Annual Review of Cell and Developmental Biology 28(1): 489–521. http://www.annualreviews.org/doi/10.1146/annurev-cellbio-092910-154055.

Piotrowski, Jeff S. et al. 2015. “Plant-Derived Antifungal Agent Poacic Acid Targets β-1,3-Glucan.” Proceedings of the National Academy of Sciences of the United States of America 112(12): E1490–97.

Ranjan, Ashish et al. 2019. “ Resistance against Sclerotinia Sclerotiorum in Soybean Involves a Reprogramming of the Phenylpropanoid Pathway and Up-Regulation of Antifungal Activity Targeting Ergosterol Biosynthesis.” Plant Biotechnology Journal: 1–15.

Rollins, Jeffrey a. 2003. “The Sclerotinia Sclerotiorum Pac1 Gene Is Required for Sclerotial Development and Virulence.” Molecular plant-microbe interactions: MPMI 16(9): 785–95.

Saitoh, Yoshimoto, Kosuke Izumitsu, Atsushi Morita, Kiminori Shimizu, et al. 2010. “ChMCO1 of Cochliobolus Heterostrophus Is a New Class of Metallo-Oxidase, Playing an Important Role in DHN-Melanization.” Mycoscience 51(5): 327–36.

Saitoh, Yoshimoto, Kosuke Izumitsu, Atsushi Morita, and Chihiro Tanaka. 2010. “A Copper-Transporting ATPase BcCCC2 Is Necessary for Pathogenicity of Botrytis Cinerea.” Molecular Genetics and Genomics 284(1): 33–43.

Schneider, Caroline A., Wayne S. Rasband, and Kevin W. Eliceiri. 2012. “NIH Image to ImageJ: 25 Years of Image Analysis.” Nature Methods 9(7): 671–75.

Schouten, Alexander et al. 2002. “Resveratrol Acts as a Natural Profungicide and Induces Self-Intoxication by a Specific Laccase.” Molecular Microbiology 43(4): 883–94.

Schouten, Alexander et al. 2008. “Involvement of the ABC Transporter BcAtrB and the Laccase BcLCC2 in Defence of Botrytis Cinerea against the Broad-Spectrum Antibiotic 2,4-Diacetylphloroglucinol.” Environmental Microbiology 10(5): 1145–57.

Schumacher, Julia. 2016. “DHN Melanin Biosynthesis in the Plant Pathogenic Fungus Botrytis Cinerea Is Based on Two Developmentally Regulated Key Enzyme (PKS)-Encoding Genes.” Molecular Microbiology 99(4): 729–48.

Szewczyk, Edyta et al. 2006. “Fusion PCR and Gene Targeting in Aspergillus Nidulans.” Nature Protocols 1(6): 3111–20.

Upadhyay, Srijana et al. 2016. “Subcellular Compartmentalization and Trafficking of the Biosynthetic Machinery for Fungal Melanin Report Subcellular Compartmentalization and Trafficking of the Biosynthetic Machinery for Fungal Melanin.” CellReports: 1–8. http://dx.doi.org/10.1016/j.celrep.2016.02.059.

Wei, Yunxie et al. 2017. “The Laccase Gene (LAC1) Is Essential for Colletotrichum Gloeosporioides Development and Virulence on Mango Leaves and Fruits.” Physiological and Molecular Plant Pathology 99: 55–64. http://dx.doi.org/10.1016/j.pmpp.2017.03.005.

Westrick, Nathaniel M. et al. 2019. “Gene Regulation of Sclerotinia Sclerotiorum during Infection of Glycine Max: On the Road to Pathogenesis.” BMC Genomics 20(1).

Wu, Jian et al. 2021. “Host-Induced Gene Silencing of Multiple Pathogenic Factors of Sclerotinia Sclerotiorum Confers Resistance to Sclerotinia Rot in Brassica Napus.” Crop Journal. https://doi.org/10.1016/j.cj.2021.08.007.

Xu, Liangsheng, Meichun Xiang, David White, and Weidong Chen. 2015. “PH Dependency of Sclerotial Development and Pathogenicity Revealed by Using Genetically Defined Oxalate-Minus Mutants of Sclerotinia Sclerotiorum.” Environmental Microbiology 17(8): 2896– 2909.

Zhang, Chunquan, Jeffrey D Bradshaw, Steven A Whitham, and John H Hill. 2010. “The Development of an Efficient Multipurpose Bean Pod Mottle Virus Viral Vector Set for Foreign Gene Expression and RNA Silencing.” Plant Physiology 153(May): 52–65.

